# ML-MAGES: A machine learning framework for multivariate genetic association analyses with genes and effect size shrinkage

**DOI:** 10.1101/2025.02.11.637655

**Authors:** Xiran Liu, Lorin Crawford, Sohini Ramachandran

## Abstract

A fundamental goal of genetics is to identify which and how genetic variants are associated with a trait, often using the regression results from genome-wide association (GWA) studies. Important methodological challenges are accounting for inflation in GWA effect estimates as well as investigating more than one trait simultaneously. We leverage machine learning approaches for these two challenges, developing a computationally efficient method called *ML-MAGES*. First, we shrink the inflation in GWA effect sizes caused by non-independence among variants using neural networks. We then cluster variant associations among multiple traits via variational inference. We compare the performance of shrinkage via neural networks to regularized regression and fine-mapping, two approaches used for addressing inflated effects but dealing with variants in focal regions of different sizes. Our neural network shrinkage outperforms both methods in approximating the true effect sizes in simulated data. Our infinite mixture clustering approach offers a flexible, data-driven way to distinguish different types of associations—trait-specific, shared across traits, or spurious—among multiple traits based on their regularized effects. Clustering applied to our neural network shrinkage results also produces consistently higher precision and recall for distinguishing gene-level associations in simulations. We demonstrate the application of *ML-MAGES* on association analyses of two quantitative traits and two binary traits in the UK Biobank (genetic and phenotypic data from 500,000 residents of the UK). Our identified associated genes from single-trait enrichment tests overlap with those having known relevant biological processes to the traits. Besides trait-specific associations, *ML-MAGES* identifies several variants with shared multi-trait associations, suggesting putative shared genetic architecture.

## 1 Introduction

Genome-wide association (GWA) studies analyze genome-wide genotype data from a large group of unrelated individuals to identify genetic mutations, usually single nucleotide polymorphisms (SNPs), that are associated with some trait such as height, molecular biomarkers, or diseases. With advancements in sequencing technologies and the emergence of biobank datasets that sample hundreds of thousands of individuals, GWA studies have proven powerful in detecting potential causal variants for various traits and diseases. Over 45,000 GWA studies investigating more than 5,000 human phenotypes have been published since 2005 [10,40].

The fundamental idea behind GWA studies is based on fitting a linear regression where each SNP genotype is treated as an independent variable and the trait value of interest is the dependent variable. The regression coefficients are “effect sizes” of variants on the trait, and their statistical significance is determined by p-values testing whether the effect of any given variant is significantly different from zero. GWA studies generally focus on a single trait of interest. Combining signals from multiple variants can increase statistical power; therefore, it is common to perform gene-level association analysis based on SNP-level GWA results of variants in each gene [16,27,46].

One major challenge lies in controlling the inflation of effect sizes, which complicates distinguishing truly associated variants from non-associated ones. Most GWA models assume genetic variants to be independent, but there are consistent non-random associations among genotypes in a sample known as linkage disequilibrium (LD). Closer mutations exhibit higher LD (i.e., correlation), which is also affected by genetic recombination rates and population history. A non-causal variant may show large statistical effect for a trait because of its linkage to a causal variant; these spurious associations make localizing truly associated variants challenging. In a region with extremely dense variants, multiple highly correlated variants may show even stronger effects than the truly causal one. Additionally, many complex traits involve thousands of contributing variants (“polygenic traits”), further complicating GWA studies [9,33,8,53].

There is a need to explore multiple traits simultaneously. Capturing complex interactions among genetic variants and traits provides a deeper understanding of associated biological pathways and mechanisms. As reviewed by Solovieff [41], current multi-trait approaches include developing models to test the association between a genetic variant and multiple traits simultaneously—often combining existing univariate GWA results to improve the power of detecting genetic associations for each trait. Methods based on univariate results focus on testing whether the variant is significantly associated with some traits [18,7] and estimating trait-specific effects [49,51]. Researchers are particularly interested in investigating variants that are trait-specific versus pleiotropic, where a single genetic variant influences multiple traits [43,54], to generate targeted hypotheses for precision medicine and drug discovery [41].

In this study, we develop a new scalable method, *ML-MAGES* (Machine Learning for Multivariate Association analyses with Genes and Effect size Shrinkage), to analyze multi-trait GWA effects using neural networks and variational inference.

### 1.1 Related methods

Estimating effects so that associated variants have non-zero effects and non-associated ones have (closeto-)zero effects is a goal for many polygenic modeling applications. Regularized regressions and Bayesian approaches have both been widely adopted to tackle this problem by introducing sparsity into the effects [57,15,29,21,61,60]. Several existing Bayesian methods have used Gaussian mixture models to introduce sparsity (i.e., zeros) into effect sizes [29,21,61,47,44,62,59,51,23] (see Appendix A.3). The underlying idea is that associated variants will exhibit non-zero effects.

A popular task in genetic research that is similar to shrinkage—but on a local scale—is fine-mapping. It aims to locate a few variants that very likely contain a causal one within some small trait-associated region obtained from a GWA study, usually by assigning posterior probabilities of causality to each variant [42,33,37] (also see Appendix A.3). A state-of-the-art fine-mapping method is *SuSiE* (its extension *SuSiE-RSS* uses only summary-level data) [56,64], which formulates the task as a variable selection problem. Although fine-mapping techniques are computationally expensive and cannot scale genome-wide, by decomposing the genome-wide task into tasks on chromosomal segments, we are able apply *SuSiE-RSS* and assess its performance against various effect size shrinkage methods.

Cheng *et al*. [14] previously developed *gene-ϵ* to categorize association effects along the genome for a single trait, distinguishing truly associated variants from spurious ones. It first accounts for inflation in the effect sizes via SNP-level shrinkage using elastic net regularization [63], then assigns variants into groups via clustering using a univariate *K*-mixture model. The variances of the zero-mean Gaussian components are used to formulate a SNP-level null threshold. The null hypothesis is then used for identifying associated genes for the trait.

## 2 Methods

Inspired by the single-trait *gene-ϵ* framework, we propose a new method, *ML-MAGES*, that uses machine learning methods to 1) effectively account for inflation in univariate GWA effects and 2) advance multivariate GWA analyses. *ML-MAGES* incorporates several advancements compared to related methods, with two key innovations: 1) use of deep learning to shrink effect sizes efficiently, and 2) use of variational inference to perform clustering for multivariate associations flexibly.

There are three main steps in *ML-MAGES* (Fig. 1): 1) effect size shrinkage, which takes in GWA summary statistics and linkage disequilibrium (LD) and gives regularized effects; 2) association clustering, which groups the variants according to how strongly they are associated to the trait(s) of interest using output from step 1; and 3) gene-level analysis, which aggregates the association clustering of variants in each gene from step 2 and differentiates whether and how genes are associated to the trait(s). We discuss each step in detail below. *ML-MAGES* takes in observed effects 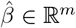 and standard errors *s* ∈ℝ^*m*^ for GWA results on *m* variants, as well as LD between pairs of variant stored in a *m × m* matrix *R*. The LD score of variant *j*, ℓ_*j*_, which quantifies the amount of genetic variation tagged by it, is the sum of its squared correlations with all other SNPs. Calculation of these input can be found in Appendix A.2.

**Fig. 1.**
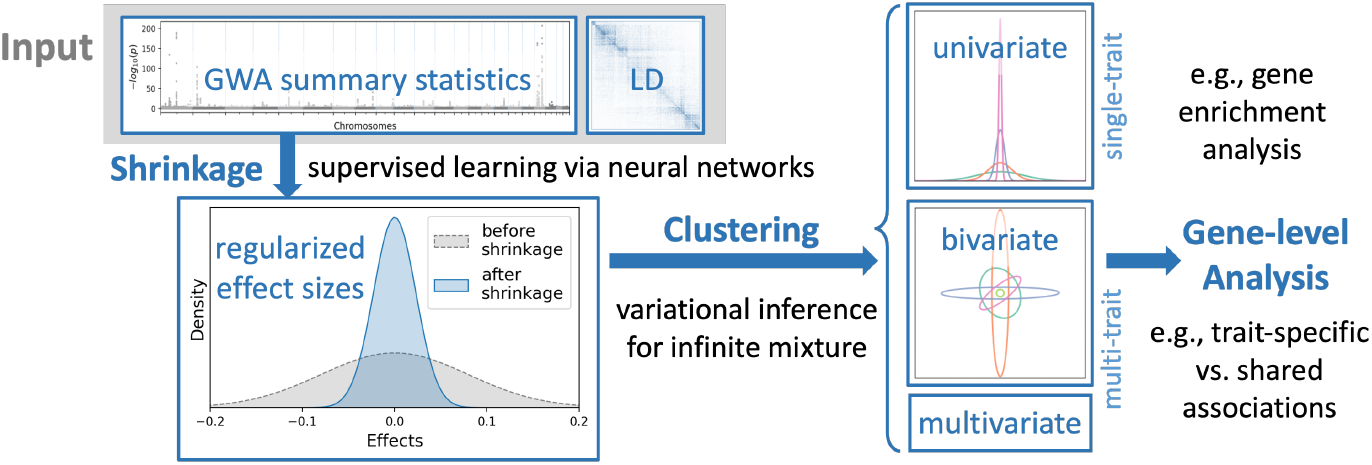
Workflow of *ML-MAGES* for assessing the associations of variants to (multiple) traits.

### 2.1 Effect size shrinkage

The genotypes of many genetic variants, especially those that are physically close together, are not independent due to linkage disequilibrium, introducing inflation into the observed effects 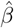 compared to the true ones *β*. The goal of effect size shrinkage is to obtain regularized effects 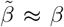 by accounting for inflation caused by correlations. The number of truly causal variants—those with 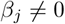—is usually limited even for highly-polygenic trait. Because of this, the goal of shrinkage is to encourage sparsity in 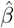 by introducing (near-)zeros so that 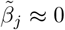 for non-associated SNPs. This makes effect size shrinkage similar to variable selection in high-dimensional data. Regularized regression is a popular choice for implementation of shrinkage. *gene-ϵ* [14] used elastic net [63] for shrinkage after testing multiple approaches (see Appendix A.3).

#### Shrinkage as supervised learning: ML-MAGES

Due to the inefficiency of regularized regression approaches and their limitations in handling non-linearities, we approach effect size shrinkage as a supervised machine learning task using feed-forward neural networks (NNs). The supervised learning formulation uses summary statistics and LD to construct feature input, denoted as Ω, and true effects *β* as the targeted output. The objective of the training problem (using the mean squared error loss) is

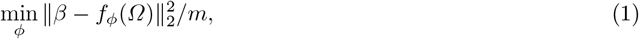

where *f*_*ϕ*_(*·*) is the function represented by NN, *ϕ* is the set of network parameters to optimize over, and *m* is the data size. With trained model parameters *ϕ*^*^, regularized effects are 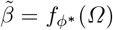. Since we need ground truth training data for supervised learning, which is not available for genetic studies, we simulate synthetic association data for training (details in Appendix A.4). Specifically, we draw true non-zero effects from a normal distribution and simulate synthetic trait values based on an additive model as in [14], from which we obtain GWA effects and apply transformations to match the observed effect size distributions.

We benchmark our NN methods against elastic net [14] and the fine-mapping method *SuSiE-RSS* [56,64] for shrinkage, both computationally expensive when applied on a large number of variants. We therefore decompose the LD matrix of each chromosome into approximately independent LD blocks and perform shrinkage on each block separately (see Appendix A.5).

#### Feature construction

We hypothesize that variants highly correlated with a focal variant contribute the most to its inflation. We therefore obtain input features for a focal variant based on the summary data of the top *T* variants in highest LD to it, with *T* ≪ *m*. The features for variant *j* are

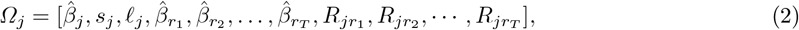

where 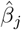 is the observed effect, *s*_*j*_ is the standard error, ℓ_*j*_ is the LD score of the variant *j*, and *r*_1_, …, *r*_*T*_ denote the indices of the top *T* variants that are most correlated to variant *j*. The number of features for each variant under this construction is 2*T* + 3. This feature construction relieves the computational burden of *ML-MAGES* by constraining the input space to be of size *O*(*Tm*), a great reduction from that of the regularized regression approaches (*O*(*m*^2^), see Appendix A.3).

#### Neural network architectures and training

We propose two NN architectures, differing in their model complexity. The first one contains two fully connected layers besides the output layer, and the second one contains three. We also test the architectures that vary in input feature size through varying *T*. Detailed architectures as well as other training settings are in Appendix A.6.

### 2.2 Clustering of associations based on effect size distributions

Next, we are interested in clustering the non-zero variants based on distributions of their regularized effects 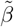 that are assumed to be zero-centered. *gene-ϵ* [14] works with the effects of a single trait, fitting a zero-mean *K*-mixture Gaussian model to classify variants as associated, non-associated with spurious signals, or non-associated with zero-effects. *ML-MAGES* generalizes this method where an arbitrary number of variant effect clusters and traits can be simultaneously considered. We use a zero-mean infinite-mixture of multivariate Gaussians for clustering, which can flexibly infer the number of variant groups. *gene-ϵ* specifies a limited range of *K* values and the optimal *K* reported sometimes equals the largest one [14], suggesting that a higher *K* possibly fits the data better. *ML-MAGES* overcomes this issue using infinite mixture, which is particularly beneficial for multi-trait analysis as the complexity of association relationships grows quickly with the number of traits.

#### Zero-mean infinite mixture model

Variants with zero or close-to-zero effects after shrinkage are non-associated, so to reduce the problem size of clustering, regularized effects are first thresholded to a subset of non-zero ones. Let 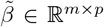 be the regularized effects of *m* variants on *p* traits. The clustering input is a matrix *γ*∈ ℝ^*J×p*^, where rows 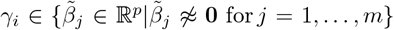 are the *J* variants with non-zero effects for any of the *p* traits. We assume that each cluster follows a zero-mean multivariate Gaussian and each variant belongs to a cluster. Let *z*_*ik*_ ∈ {0, 1} be the latent indicator variable with *z*_*ik*_ = 1 if variant *i* belongs to cluster *k* and 0 otherwise.

Let 𝒩 (*µ, A*^−1^) be a multivariate normal distribution with mean *µ* and covariance matrix *A*^−1^, or equivalently, with precision matrix *A*. Let Cat(*π*) be a categorical distribution parameterized by *π*. The zero-mean infinite-mixture model is

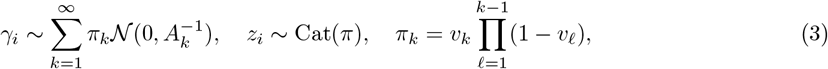

where *π*_*k*_ denotes the probability that *γ*_*i*_ is drawn from cluster *k*, with 0 *< π*_*k*_ ≤ 1 and 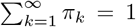.The sequence of mixture weights *π* can be derived from the stick-breaking construction of a Dirichlet process as 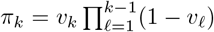 [38,6], where *v*_*k*_’s are drawn from a Beta distribution *v*_*k*_ ~ Beta(1, *α*). We approximate the model posterior and estimate parameters using variational inference (VI) [25,55], with steps detailed in Appendices A.7 and A.8.

Each inferred cluster is a zero-mean Gaussian with covariance term 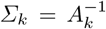.The inferred clusters are first ordered by decreasing *π*_*k*_ values, then an optimal number of clusters, *K*^*^ (*K*^*^ ≤ *K*), is determined by truncating *K* at a reasonably large value such that 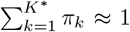.The remaining *K*^*^ clusters are then ordered decreasingly in Tr(*Σ*_*k*_). When *p* = 1, this is equivalent to the ordering 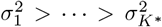 used by *gene-ϵ*. The clustering label of variant *i* is arg max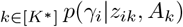.

### 2.3 Gene-level analysis

Combining results from SNPs in each gene enables gene-level association analysis. Gene-level signals can be compared with existing biological knowledge and offer mechanistic insights into the genetic pathways underlying the traits. When *p* = 1, *gene-ϵ* formulates a SNP-level null hypothesis based on variance of clusters and compute an enrichment test statistic for the gene association [14] (described in Appendix A.9). When *p >* 1, despite the lack of a single test statistic, patterns can still be extracted from the effect size distributions of variants in each cluster, both visually and quantitatively.

#### Multivariate gene-level association: visualization and summarization

When *p* = 2 or *p* = 3, clusters can be visualized as ellipses and ellipsoids. For higher *p*, a Gaussian mixture component can be summarized by the eigenvalues and eigenvectors of its variance-covariance matrix *Σ*_*k*_, which provide a geometric interpretation of the distribution. For example, if the major axis of a cluster falls close to an Cartesian axis and is much larger than all the minor axes, then the cluster can be interpreted as being “trait-specific”, with variants strongly associated to one trait but not the other. If Tr(*Σ*_*k*_) is large and angle of the major axis to any of the Cartesian axes is large, the cluster likely contains shared associations. If Tr(*Σ*_*k*_) is small, the cluster has spurious associations. We compute fractions of variants in each gene that fall into different types of clusters (see Appendix A.10). Genes with large shared-association fractions are of interest for studying putative pleiotropy.

We also calculate the normalized sum of products of SNP effects across traits in each gene as 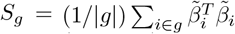, where *g* is the set of variants in a gene and 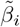 of size |*g*| *× p* are their effects. Having large (or small) values in different entries of matrix *S*_*g*_ indicate different association relationships and strengths. For example, if a gene is associated to trait *j* but not the others, then *S*_*g*_(*j, j*) would be large while *S*_*g*_(*j*^*′*^, *j*^*′*^) for *j*^*′*^≠ *j* are small. For *S*_*g*_(*j, j*^*′*^) large, the gene is likely associated to both traits *j* and *j*^*′*^, with associations positively correlated.

### 2.4 Performance comparisons

We compare the performance of neural-network-based shrinkage against the regularized regression (elastic net [14], denoted as *enet*) and the fine-mapping method (*SuSiE-RSS* [64], denoted as *susie*). We implement and compare six neural network models: the combinations of three different feature sets—with *T* =5, 10, and 15 in Eq. 2—and two architectures—the two-layer and three-layer networks. We evaluate their performance on simulation data (Appendix A.4). To demonstrate the robustness of the method, we adopt the idea of ensemble learning in additional experiments, where we train ten models of the same architecture each using a randomly sampled subset of training and validation data, and average the regularized effects from ten models to be the final outcome (see Appendix A.13).

To quantify the performance of all methods on shrinkage and gene-level analysis, we use the following measures. On SNP-level comparison, we assess how good the regularized effects 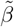 approximate the true effects *β*. We compute root mean squared error (RMSE) and Pearson correlation between them. We also assess how successfully each shrinkage method can rank truly associated effects over non-associated ones via the precision-recall curves (PRCs), with a curve closer to the upper-right corner indicating better performance. For gene-level performance comparison, we conduct single-trait gene enrichment tests, assessing the power for identifying associated genes at a significance level of *p* = 0.05, corrected for multiple testing by controlling the false discovery rates (FDR) at level *α* = 0.05 [4]. We compare the precision, recall, and F-score of identifying truly associated genes for each method. For bivariate analysis of two traits, we first obtain the fraction of variants in each gene that belong to trait-specific clusters, and rank it against the ground truth of whether each gene is simulated to have trait-specific association. Similarly, we rank the fraction of variants in clusters of shared associations against their ground truth. The rest of the clusters are categorized as spurious associations (see Appendix A.10). F-scores are reported for these comparisons, with a higher score indicating better ability to distinguish associations that are trait-specific versus shared.

## 3 Results

### 3.1 Performance comparison on simulation data

The six neural-network models show very similar performances across all the comparisons. We focus on the result for the 2-layer and 3-layer models using *T* = 15 variants for features (denoted as *ML-MAGES 2L* and *3L*, respectively). We compare them to elastic net (*enet*) [14]) and *SuSie-RSS* (*susie*) [64]). We leave an exhaustive list of comparisons of all 8 shrinkage methods to Tables 1-4 in the Appendix. The 6 NN models have also been trained (and validated) on simulation based on a different set of chromosomes, and their performances are consistent (results not shown).

#### Approximating true effects at the SNP-level

We first compare the performance of methods on effect size shrinkage. We show the SNP-level comparison results on gene-level simulation in Fig. 2A and B. Similar results on SNP-only simulation are included in the Appendix (Table 1). The precision-recall curve (PRC) of *susie* shows comparable performance to *ML-MAGES*, all better than *enet* (Fig. 2A), suggesting that methods are ranking truly associated variants highly. Additionally, our NN methods outperform *enet* and *susie* in RMSE (Fig. 2B). We note that the PRC shows a sharp decrease in precision as recall becomes larger. This pattern occurs when data evaluated is highly imbalanced, which is indeed the case in our scenario as approximately only 1.5% of variants are simulated to be truly causal. Recall that as a fine-mapping method, *susie* is designed for selecting a few variants that very likely contain the true causal ones from a small region, which is exactly what the gene-level simulation is generating. However, *susie* aims to locate the few causal variants, therefore tending to overestimate their effect sizes, giving a less desirable RMSE. A good approximation of the true effects is important for accurate clustering in the next step. Our NN-based method achieves the highest accuracy in this task, and therefore is a competitive approach for effect size shrinkage. Also note that training is based on SNP-only simulations where associated variants are not densely distributed within small segments, which does not quite assume the same generative process as the gene-level simulation, but the NN models still perform well on the task.

**Fig. 2.**
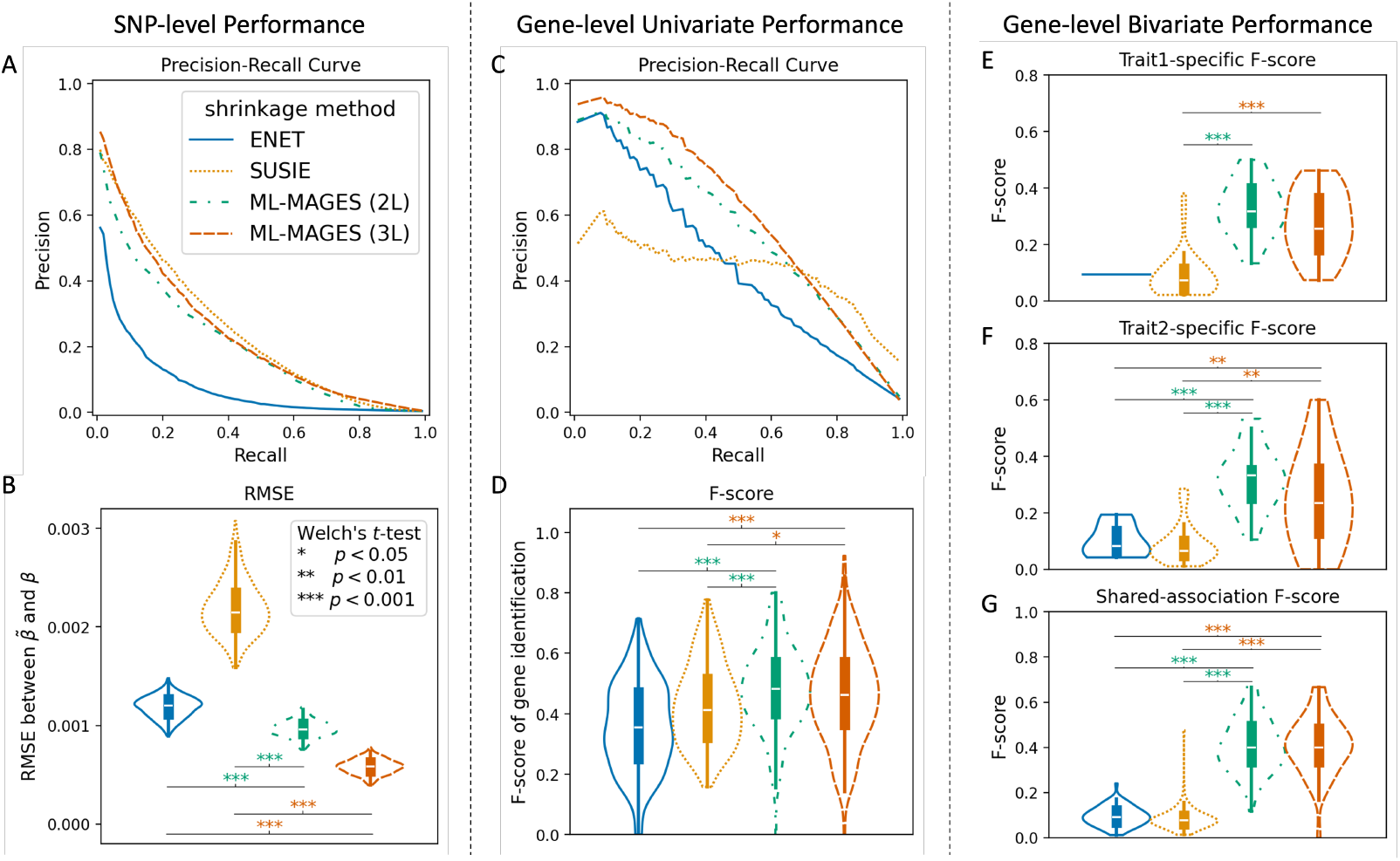
Our NN methods *ML-MAGES 2L* and *3L* outperform *enet* and *susie* on shrinking inflated GWA effect sizes in simulations. Left column: SNP-level performance, evaluated by comparing the regularized effects and the true effects of each simulation. **Middle column**: gene-level single-trait analysis performance, evaluated by comparing univariate enrichment test with the simulated ground truth. **Right column**: gene-level two-trait analysis performance, evaluated by comparing bivariate clustering output with the simulated ground truth. Legends shown in panel A apply to all panels; each violin plot ordered from left to right as *enet, susie*, and *ML-MAGES 2L* and *3L*. Violin plots are labeled with the significance level of Welch’s *t*-test for difference between measures of our methods and the other two, as shown in panel B legends. **A**: Precision-recall curve (PRC) averaged across all 200 simulations (by interpolation), where the positives are the true non-zero effects and the precision-recall pairs are obtained by thresholding 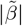. **B**: RMSE between *β* and 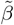. **C**: PRC of each method averaged across all simulations (by interpolation), where the true positives are the true associated genes and the precision-recall pairs are obtained for different p-values of gene enrichment tests. **D**: F-score of identifying true associated genes, where genes with a FDR-adjusted *p <* 0.05 from the enrichment analysis is identified as associated. **E** and **F**: Trait-specific F-scores for identifying genes only associated to one simulated trait, when ranking the genes by their fraction of variants in trait-specific clusters and compared against being truly trait-specific, for simulated trait 1 and 2. **G**: F-score for identifying genes associated to both traits, when ranking the genes by their fraction of variants in shared association and compared against being truly associated to both traits.

Another strength of our NN shrinkage is its computational efficiency. We compare the time taken by different methods to shrink the same set of simulated effects—based on genotyped data of 15,250 variants on chromosome 15 from UK Biobank [45]. Our NN methods (*ML-MAGES* : 0.86s) are more than 10x faster compared to *enet* (15.88s) and *susie* (33.62s) when averaged across 200 simulations (see Appendix Table 5). Even considering the one-time training that takes 15 to 25 seconds per epoch for less than a hundred epochs on our simulation data (Appendix Table 6), using NN models is more efficient. Moreover, the computational burden of NN models will not scale with the number of variants exponentially as in *enet* or *susie*, which poses challenges on analyzing larger sets of variants (e.g., the imputed whole-genome sequence data).

#### Distinguishing genes with different types of associations

Besides SNP-level performance, we also compare the gene-enrichment analysis results based on each method’s shrinkage output and subsequent clustering. We assess the power for identifying associated genes at a significance level of *p* = 0.05 (FDR-corrected). We show the PRCs and F-scores of identifying truly associated genes correctly (Fig. 2C and D). *enet* and *susie*-based results perform poorly on both measures, which is expected based on their SNP-level performance (Fig. 2B). *susie* shows a high recall but a low precision with many false positives (Appendix Table 3), which is not surprising since it is not designed to identify causal genes. We also evaluate the performance of these methods on multi-trait analysis, using the simulated bivariate data as an illustrative example. We compute the F-scores of gene ratings according to fraction of variants in each type of cluster against their true association type (Fig. 2E-G). These scores indicate the ability of the methods to identify trait-specific genes versus those with shared associations. *enet* has only one simulation that identifies a trait-1-specific cluster, resulting in a single bar in Fig. 2E. *ML-MAGES* performs well for the gene-level analyses.

To further showcase the strength of multi-trait analysis, we included a simulation of three traits which captures more complex shared association scenarios in Appendix A.12 and Fig. 7. The simulation procedure mirrors the bivariate one. The *ML-MAGES* framework consistently outperforms elastic net in identifying truly associated SNPs and genes, as well as detecting the correct shared association patterns among simulated traits, as shown by PRCs and F-scores in Fig. 7.

#### Comparing performance when models are trained on different simulation settings

In our previous simulation analyses, the synthetic training data was generated using the same type of real data as the evaluation data. To demonstrate the reliability of *ML-MAGES* when this does not necessarily hold, we perform additional simulation analyses. We show the performance of models trained for a more challenging shrinkage task when the simulated training data are based on different data and settings from the intended evaluation set. Specifically, we simulate training data based on imputation data in UKB with genetic markers not directly genotyped, which is more than eight times larger than the genotyped array data size. The much higher density of the imputed variants causes significantly larger LD between them, which subsequently introduces large inflation in the effects, making the shrinkage problem much more challenging than in the genotyped data. Although the model’s training performance is less optimal when using simulations based on imputed data, models trained on imputed data still consistently outperform the elastic net method in shrinkage performance when applied to simulations based on genotyped data. This holds across various simulation settings that differ in the proportion of true causal genes and causal variants. We detail the simulations and comparison of SNP-level performance (Fig. 8) in Appendix A.13.

#### Comparing to a linear model

To highlight the advantage of *ML-MAGES* ‘s ability to capture nonlinear relationships among inflated effects, we compare its performance to a single-layer neural network within the same framework—an architecture limited to modeling linearities. As illustrated in Fig. 8 (Appendix A.13), the results demonstrate the benefit of accounting for nonlinearities in effect size shrinkage. The linear model consistently underperforms nonlinear ones in estimating the true effect sizes of associated variants.

### 3.2 Real data analysis

#### Applying the method to the GWA of two continuous traits: HDL and LDL

To get a more comprehensive understanding of how the regularized effects from different shrinkage methods affect gene-level enrichment tests, we apply all methods on GWA effects of two quantitative traits from UK Biobank [45]: high-density lipoprotein cholesterol (HDL) and low-density lipoprotein cholesterol (LDL). The data contains GWA results on 489,953 variants from 334,851 European-ancestry individuals (see Appendix A.1 for a description of the data). All results shown in Fig. 3 except for panel C and D are based on *ML-MAGES (3L)*.

**Fig. 3.**
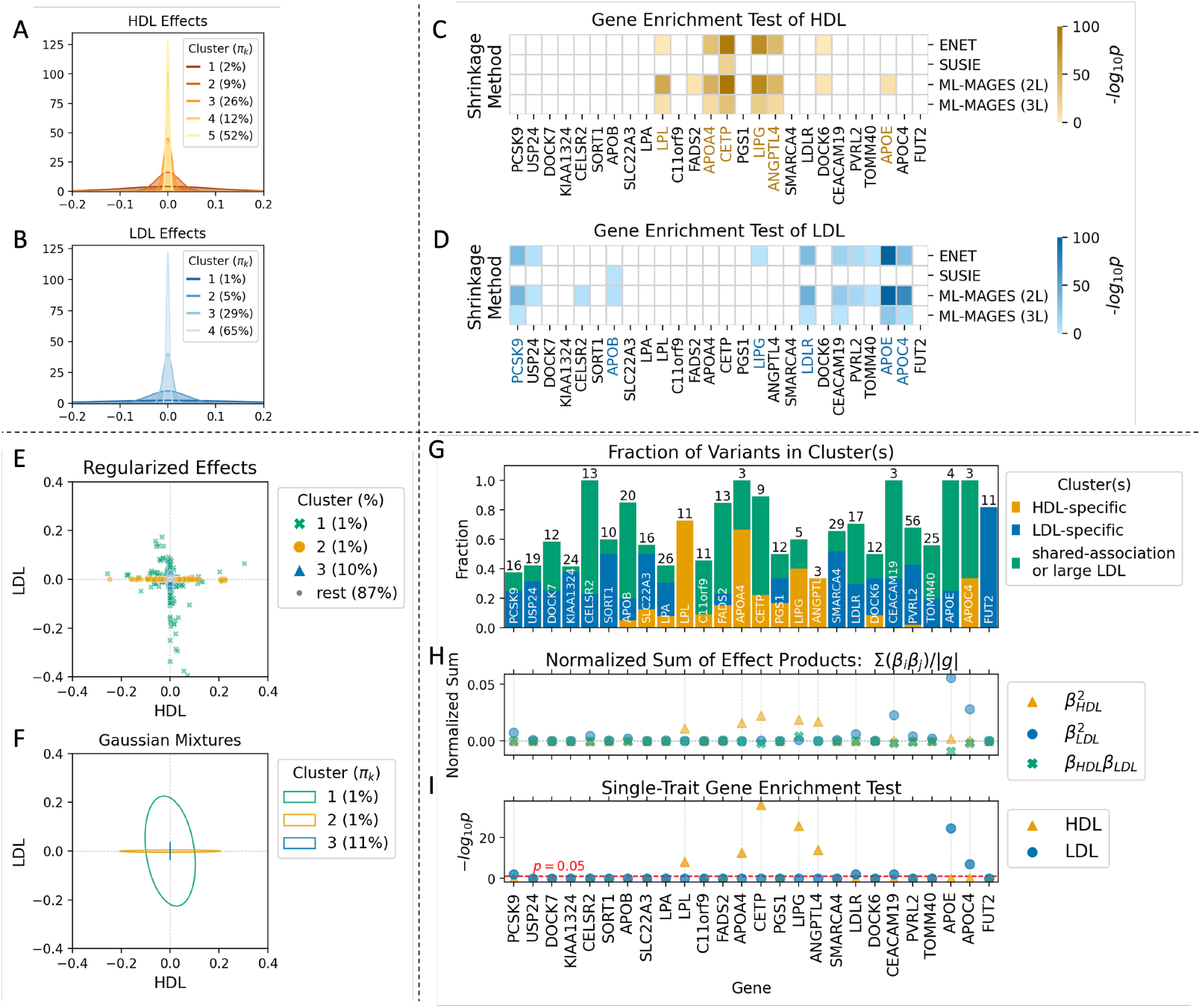
Clustering and gene-level analysis results, using shrinkage output on GWA effects of traits highdensity lipoprotein cholesterol (HDL) and low-density lipoprotein cholesterol (LDL) in UK Biobank European individuals. For panels C and D, the results using 4 shrinkage methods (*enet, susie*, and *ML-MAGES 2L* and *3L*, see also Fig. 2) are all presented. For the other panels, only the results of *3L* are included. **A** and **B**: The univariate clustering on GWA effects of HDL and LDL. Clusters are represented by the corresponding Gaussian 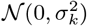, labeled with their inferred mixing weights *π*_*k*_. **C** and **D**: The enrichment analysis of genes associated to HDL and LDL. The set of genes shown are those included in panel G. The cells are colored by − log_10_(*p*) values of the test, with darker color indicating higher statistical significance, and non-significant genes with adjusted *p*≥ 0.05 are colored white. Associated genes that have related biological processes terms in the GO Biological Processes network [3] are highlighted in color. Unlike the other three methods, *susie* shrinkage fails to identify the most relevant genes. **E**: The bivariate clustering results based on the regularized effects of HDL and LDL. Clusters are color-coded and variants are styled differently by their cluster labels. Clusters are ordered in descending Tr(*Σ*_*k*_). The proportion of variants with non-zero effects in each cluster is labeled in the legend. Clusters beyond the first three are categorized as spurious associations and grouped into the “rest”. **F**: Inferred mixtures from bivariate clustering, shown as a scaled confidence ellipse for each corresponding Gaussian mixture 𝒩 (0, *Σ*_*k*_), with inferred mixing weights *π*_*k*_ labeled in the legend. **G**: The fraction of variants assigned to each of the listed genes that belong to each type of associated clusters. The genes listed either have at least 10 variants and more than 40% of variants from one of the non-spurious clusters, or are identified by enrichment test of either of the two traits (panels C and D). The number of variants assigned to each gene is annotated on the top of the bars. **H**: Normalized sum of squared effects or sum of effect products of variants in each gene, calculated as 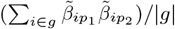,where *g* denotes the set of variants assigned to a gene with variants indexed by *i*, and *p*_1_ and *p*_2_ index the two traits. Three sums are considered: two trait-specific sum of squared effects, and the sum of effect products for shared associations. Genes having large trait-specific sums are those identified by the enrichment test (marked by dashed lines). **I**: Enrichment analysis − log_10_(*p*) for each of the two traits, corresponding to the color-coded values in bottom rows of panels C and D. Genes identified by the enrichment test are marked by vertical dashed lines.

Genes related to biological processes of HDL and LDL have been widely-studied and the information is available from the GO knowledge base [3]. We perform gene enrichment analysis on each of the two traits separately using their regularized effects and clustering output. We can therefore verify the top associated genes identified with their known biological processes. The enrichment test results based on all methods are shown in Fig. 3C and D. *enet* and two NN methods rank a very similar set of genes the highest, with majority of them showing related biological processes, while *susie* gives an almost entirely different set of genes that mostly do not appear in the knowledge base. The poor performance of *susie* is consistent with our observations on simulation (Fig. 2). We should point out that although not doing well on simulation, *enet* shows comparable output to the NN methods in this application, likely because it manages to keep at least some variants in the strongly associated genes with non-zero effects. This also explains why *enet* performs well in the work of [14].

For multi-trait analysis, we are able to identify potentially associated genes that are trait-specific and those that are putatively pleiotropic. We refer to Fig. 3E-I for a visualization of the results. It can be clearly observed from the inferred Gaussians (Fig. 3F) that some variants have trait-specific associations (e.g., clusters 2 and 3) and some have shared associations (e.g., cluster 1). The Gaussians with the smallest Tr(*Σ*_*k*_) (e.g., clusters labeled “rest”) are those that contain likely spurious associations, similar to the univariate scenario in *gene-ϵ*. In the bivariate analysis, we group clusters into three types based on their *Σ*_*k*_’s (Appendix A.10): specific to HDL, specific to LDL, and associated to both or having large effect in LDL. The fraction of variants in each gene that belong to each type of clusters (Fig. 3G) suggests some genes being specifically associated to one of the traits while some others having shared associations. We subset genes with strongest signals for visualization. Genes identified by the univariate enrichment test (Fig. 3I) correspond to those with both large trait-specific fractions and a large number of variants. The enrichment analysis only uses a SNP-level null threshold obtained from the clustering results, potentially missing out some weakly-associated genes and not fully utilizing SNP-level information. Instead, our multi-trait analysis is able to bring more information from the SNP-level effects to the gene level (Fig. 3G and H) while capturing signals for shared associations and help locate genes potentially of interest for the study of their trait-specific versus pleiotropic contributions.

#### Demonstrating the method in application to two binary traits

To demonstrate the generalizability of *ML-MAGES*, we apply the models trained using imputation data—as described in the previous section—onto the GWA results of two binary traits in the UKB data, restricted to European ancestry individuals. Both traits are ICD10-coded diseases: C50 (malignant neoplasm of breast) and Z90.1 (acquired absence of breast). The results are shown in Fig. 4. The log odds ratios of logistic regression from binary GWA are encoded as effects of the variants. *ML-MAGES* identifies two most significant genes that have shared associations with both diseases, *FGFR2* and *TOX3*, suggesting a likely similarity in their biological underpinnings. These two genes show the largest normalized sum of effect products, and they exhibit very similar patterns in the fraction of variants belonging to each cluster (Fig. 4). These two genes were previously reported by Cortes *et al*. [17] to be associated with nearly identical nodes in the ICD-10 ontology. The identified variants pointed out by Cortes *et al*. [17] that localize them are rs2981575 and rs4784227, respectively, and both variants are in the cluster denoting shared associations in *ML-MAGES* ‘s clustering. This further supports the validity of our method in multi-trait association analysis.

**Fig. 4.**
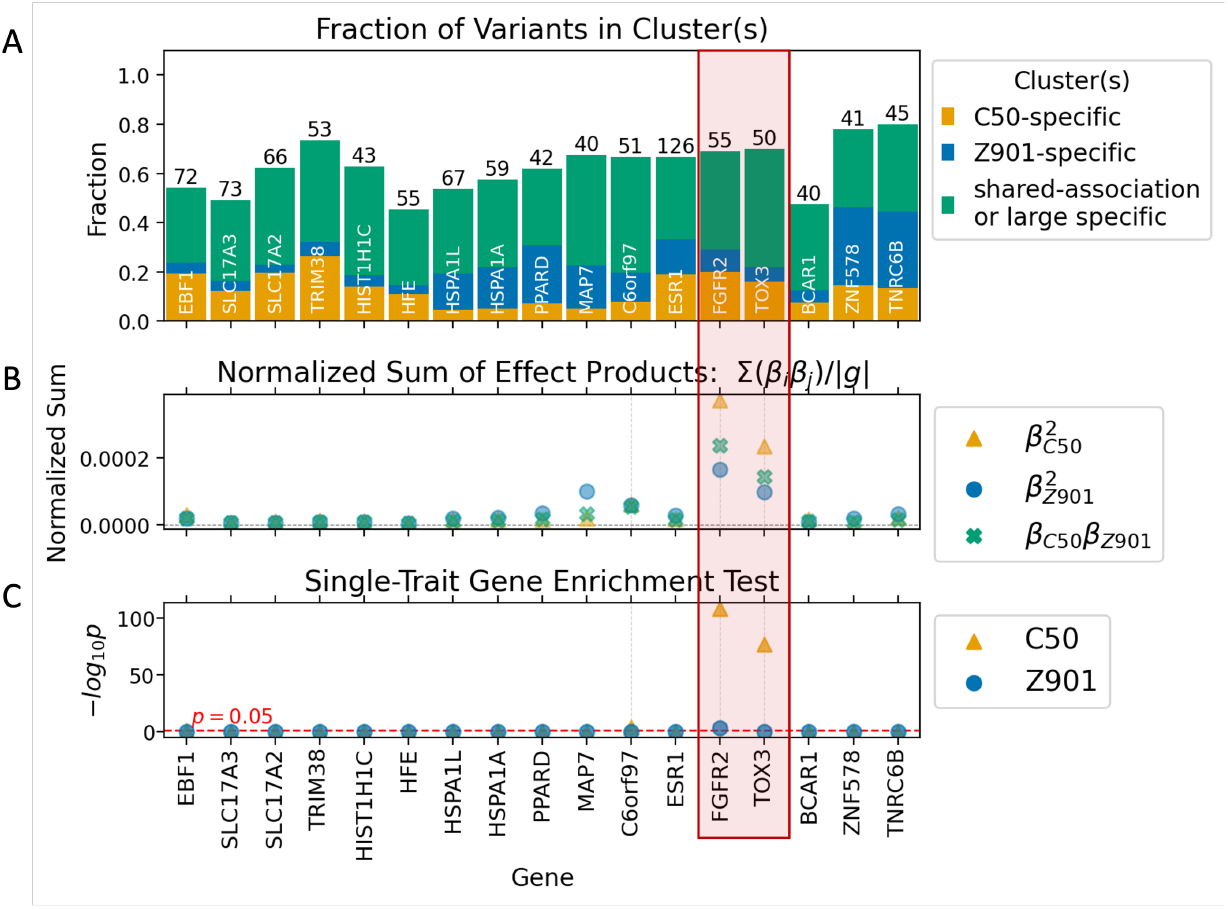
Gene-level analysis results, using shrinkage output on GWA effects of two binary traits in UK Biobank European individuals. Two binary traits are C50 (malignant neoplasm of breast) and Z90.1 (acquired absence of breast). The figure style follows that of Fig. 3 panels G-I. **A**: The fraction of variants in each gene that belong to each type of clusters. The genes listed have at least 40 variants and more than 30% of variants from one of the non-spurious clusters. **B**: Normalized sum of effect products of variants in each gene. **C**: Enrichment analysis − log_10_(*p*) for each trait. Genes identified by the enrichment test are marked by vertical dashed lines. The two genes in the red box, FGFR2 and TOX3, are those identified by Cortes *et al*. [17] as showing similar biological pathways.

## 4 Discussion

Genome-wide association (GWA) studies have been powerful for generating hypotheses regarding associated genetic variants and the genetic architectures of numerous complex traits in humans [52,50]. We introduce *ML-MAGES*, which uses machine learning approaches to perform multi-trait association analyses, addressing two open challenges: inflation in GWA effects caused by LD, and shared associations across traits undetectable by single-trait analyses. Our method is a computationally efficient and functionally effective tool to tackle these challenges.

Our neural-network based shrinkage achieves higher accuracy in approximating the true SNP-level effect sizes than do regularized regression [14] and fine-mapping [64] methods (Fig. 2A and B) while improving the computational efficiency (Appendix Table 5), and in turn shows better performance in categorizing gene-level signals in the simulated data (Fig. 2C-G). Our generalization of *gene-ϵ*’s univariate clustering [14] to the multivariate scenario enables exploration of multi-trait association relationships beyond estimating the single-trait SNP-level null thresholds. Moreover, our infinite mixture model for clustering allows an optimal number of clusters to be inferred from data, overcoming the potential underestimation of the complexity of effect size distributions in a fixed *K*-mixture model. Aggregating SNP-level results at the gene-level, our new method presents rich information regarding how variants and genes show trait-specific versus shared associations across-trait. It provides summary values (e.g., angles and ratios of axes for Gaussian ellipses, fractions of variants in each clusters, Fig. 3F and G) that can be interpreted both quantitatively and visually for an intuitive understanding of signals in multi-trait associations.

We acknowledge that a major drawback of using supervised learning for shrinkage is its reliance on simulated data for model training. Generation of the simulation data requires individual-level genotypes and synthetic GWA effects need to be similarly distributed as real ones. These impose limitations on the quality of the models when simulated data are used for different organism, or even different populations—for example, simulations based on human genome will likely not be applicable to plant genetics. In this study, we focus our analysis on European-ancestry individuals. In the context of GWA in populations with different or admixed ancestries, models trained on European-based data will show a decrease in their performances, which is a common observation in genetic studies [34,39,30]. Moreover, the distribution of real effects may not always be centered at zero, in which case the quality of the model may be further compromised. To partially overcome this issue, besides the symmetric Laplace transformation we used in the preparation of training data, we also include an asymmetric option to accommodate the case when effects are not symmetric around zero. A downside compared of our approach to methods like elastic net shrinkage is that we require generating synthetic data. However, additional simulation analyses show that supervised learning-based shrinkage outperforms elastic net shrinkage even when the training and evaluation simulations are generated using different real data, given that the effect size distributions are matched (Appendix Fig. 8). Nevertheless, generating realistic synthetic data adaptively continues to be a challenge in genetic studies, and further advancement in simulation strategies would benefit our method as well.

Another limitation of our method is the difficulty in interpreting results of multivariate clustering. As observed from our demonstrative example, a locally-optimal cluster sometimes groups variants with different types of effects together—for example, some being putatively pleiotropic and some having large trait-specific effects (Cluster 1 in Fig. 2E). The inferred Gaussian would instead suggest shared associations strong for one trait and weak for the other (Fig. 2F). We need to be careful when designating the association type of the clusters—visualization (like Fig. 2E and F) can help detect these patterns, but not always feasible. Visualizing clusters becomes particularly challenging when analyzing more than two or three traits. Methods to intuitively summarize and present the clustering results, especially for high-dimensional data, would be beneficial. Formulating multivariate null hypotheses can also help. That is, to come up with “null thresholds” for multi-trait association types and their corresponding “enrichment tests” to assess whether each gene shows specific association. Other analyses to examine the variants fall under different clusters are possible. For example, calculating a score similar to the polygenic score for each individual by adding up their genotypes weighted by corresponding effects on sets of variants in each cluster and comparing the scores to the trait values can implicate genetic liability of the traits to variants in different clusters. Estimating the heritability contributed by SNPs in each cluster can also help quantify their informativeness.

Multi-trait association analyses can help uncover potential pleiotropic effects and contribute to the development of novel therapeutic hypotheses regarding multiple diseases or medical conditions. Given the intricate genetic architecture underlying complex traits, analyzing multiple traits simultaneously offers distinct over examining each trait individually. It allows for the identification of shared genetic associations across some or all traits within a set, providing a more comprehensive understanding of the genetic factors influencing them—whether or not the traits have similar characteristics. Our multivariate approach performs comparably to univariate ones for trait-specific associations yet reveals additional patterns when correlations between traits exist. Beyond multi-trait analysis, *ML-MAGES* is also applicable to the study of group-specific versus shared associations across groups like different sexes and disease onsets. With the improved efficiency and expanded functionality enabled by machine learning, our method is a complementary advance beyond single-trait GWA analyses.

## Acknowledgments

This study was funded by the US National Institutes of Health (NIH R35 GM139628) and the Data Science Institute at Brown University. Access to UK Biobank data is under UK Biobank application 22419. We also acknowledge Samuel Pattillo Smith and Wei Cheng for helpful discussions. This research was conducted using computational resources at the Center for Computation and Visualization, Brown University.

## References

1. Abraham, G., Qiu, Y., Inouye, M.: FlashPCA2: principal component analysis of biobank-scale genotype datasets. Bioinformatics 33(17), 2776–2778 (May 2017). 10.1093/bioinformatics/btx299, http://dx.doi.org/10.1093/bioinformatics/btx299

2. Aleksander, S.A., Balhoff, J., Carbon, S., Cherry, J.M., Drabkin, H.J., Ebert, D., Feuermann, M., Gaudet, P., Harris, N.L., Hill, D.P., et al.: The gene ontology knowledgebase in 2023. GENETICS 224(1) (Mar 2023). 10.1093/genetics/iyad031, http://dx.doi.org/10.1093/genetics/iyad031

3. Ashburner, M., Ball, C.A., Blake, J.A., Botstein, D., Butler, H., Cherry, J.M., Davis, A.P., Dolinski, K., Dwight, S.S., Eppig, J.T., Harris, M.A., Hill, D.P., Issel-Tarver, L., Kasarskis, A., Lewis, S., Matese, J.C., Richardson, J.E., Ringwald, M., Rubin, G.M., Sherlock, G.: Gene ontology: tool for the unification of biology. Nature Genetics 25(1), 25–29 (May 2000). 10.1038/75556, http://dx.doi.org/10.1038/75556

4. Benjamini, Y., Hochberg, Y.: Controlling the false discovery rate: A practical and powerful approach to multiple testing. Journal of the Royal Statistical Society Series B: Statistical Methodology 57(1), 289–300 (Jan 1995). 10.1111/j.2517-6161.1995.tb02031.x, http://dx.doi.org/10.1111/j.2517-6161.1995.tb02031.x

5. Berisa, T., Pickrell, J.K.: Approximately independent linkage disequilibrium blocks in human populations. Bioinformatics 32(2), 283–285 (Sep 2015). 10.1093/bioinformatics/btv546, http://dx.doi.org/10.1093/bioinformatics/btv546

6. Blei, D.M., Jordan, M.I.: Variational inference for dirichlet process mixtures. Bayesian Analysis 1(1) (Mar 2006). 10.1214/06-ba104, http://dx.doi.org/10.1214/06-BA104

7. Bolormaa, S., Pryce, J.E., Reverter, A., Zhang, Y., Barendse, W., Kemper, K., Tier, B., Savin, K., Hayes, B.J., Goddard, M.E.: A multi-trait, meta-analysis for detecting pleiotropic polymorphisms for stature, fatness and reproduction in beef cattle. PLoS Genetics 10(3), e1004198 (Mar 2014). 10.1371/journal.pgen.1004198, http://dx.doi.org/10.1371/journal.pgen.1004198

8. Boyle, E.A., Li, Y.I., Pritchard, J.K.: An expanded view of complex traits: From polygenic to omnigenic. Cell 169(7), 1177–1186 (Jun 2017). 10.1016/j.cell.2017.05.038, http://dx.doi.org/10.1016/j.cell.2017.05.038

9. Bulik-Sullivan, B.K., Loh, P.R., Finucane, H.K., Ripke, S., Yang, J., Patterson, N., Daly, M.J., Price, A.L., Neale, B.M.: Ld score regression distinguishes confounding from polygenicity in genome-wide association studies. Nature Genetics 47(3), 291–295 (Feb 2015). 10.1038/ng.3211, http://dx.doi.org/10.1038/ng.3211

10. Buniello, A., MacArthur, J.A.L., Cerezo, M., Harris, L.W., Hayhurst, J., Malangone, C., McMahon, A., Morales, J., Mountjoy, E., Sollis, E., Suveges, D., Vrousgou, O., Whetzel, P.L., Amode, R., Guillen, J.A., Riat, H.S., Trevanion, S.J., Hall, P., Junkins, H., Flicek, P., Burdett, T., Hindorff, L.A., Cunningham, F., Parkinson, H.: The NHGRI-EBI GWAS catalog of published genome-wide association studies, targeted arrays and summary statistics 2019. Nucleic Acids Research 47(D1), D1005–D1012 (Nov 2018). 10.1093/nar/gky1120, http://dx.doi.org/10.1093/nar/gky1120

11. Cantor, R.M., Lange, K., Sinsheimer, J.S.: Prioritizing gwas results: A review of statistical methods and recommendations for their application. The American Journal of Human Genetics 86(1), 6–22 (Jan 2010). 10.1016/j.ajhg.2009.11.017, http://dx.doi.org/10.1016/j.ajhg.2009.11.017

12. Chang, C.C., Chow, C.C., Tellier, L.C., Vattikuti, S., Purcell, S.M., Lee, J.J.: Second-generation plink: rising to the challenge of larger and richer datasets. GigaScience 4(1) (Feb 2015). 10.1186/s13742-015-0047-8, http://dx.doi.org/10.1186/s13742-015-0047-8

13. Chen, E.Y., Tan, C.M., Kou, Y., Duan, Q., Wang, Z., Meirelles, G.V., Clark, N.R., Ma’ayan, A.: Enrichr: interactive and collaborative html5 gene list enrichment analysis tool. BMC Bioinformatics 14(1) (Apr 2013). 10.1186/1471-2105-14-128, http://dx.doi.org/10.1186/1471-2105-14-128

14. Cheng, W., Ramachandran, S., Crawford, L.: Estimation of non-null snp effect size distributions enables the detection of enriched genes underlying complex traits. PLOS Genetics 16(6), e1008855 (Jun 2020). 10.1371/journal.pgen.1008855, http://dx.doi.org/10.1371/journal.pgen.1008855

15. Cho, S., Kim, K., Kim, Y.J., Lee, J., Cho, Y.S., Lee, J., Han, B., Kim, H., Ott, J., Park, T.: Joint identification of multiple genetic variants via elastic-net variable selection in a genome-wide association analysis. Annals of Human Genetics 74(5), 416–428 (Aug 2010). 10.1111/j.1469-1809.2010.00597.x, http://dx.doi.org/10.1111/j.1469-1809.2010.00597.x

16. Conneely, K.N., Boehnke, M.: So many correlated tests, so little time! rapid adjustment of p values for multiple correlated tests. The American Journal of Human Genetics 81(6), 1158–1168 (Dec 2007). 10.1086/522036, http://dx.doi.org/10.1086/522036

17. Cortes, A., Albers, P.K., Dendrou, C.A., Fugger, L., McVean, G.: Identifying cross-disease components of genetic risk across hospital data in the uk biobank. Nature Genetics 52(1), 126–134 (2020)

18. Cotsapas, C., Voight, B.F., Rossin, E., Lage, K., Neale, B.M., Wallace, C., Abecasis, G.R., Barrett, J.C., Behrens, T., Cho, J., De Jager, P.L., Elder, J.T., Graham, R.R., Gregersen, P., Klareskog, L., Siminovitch, K.A., van Heel, D.A., Wijmenga, C., Worthington, J., Todd, J.A., Hafler, D.A., Rich, S.S., Daly, M.J.: Pervasive sharing of genetic effects in autoimmune disease. PLoS Genetics 7(8), e1002254 (Aug 2011). 10.1371/journal. pgen.1002254, http://dx.doi.org/10.1371/journal.pgen.1002254

19. Davies, R.B.: Algorithm as 155: The distribution of a linear combination of χ^2^ random variables. Applied Statistics 29(3), 323 (1980). 10.2307/2346911, http://dx.doi.org/10.2307/2346911

20. Fairley, S., Lowy-Gallego, E., Perry, E., Flicek, P.: The International Genome Sample Resource (IGSR) collection of open human genomic variation resources. Nucleic Acids Research 48(D1), D941–D947 (Oct 2019). 10.1093/nar/gkz836, http://dx.doi.org/10.1093/nar/gkz836

21. Guan, Y., Stephens, M.: Bayesian variable selection regression for genome-wide association studies and other large-scale problems. The Annals of Applied Statistics 5(3) (Sep 2011). 10.1214/11-aoas455, http://dx.doi.org/10.1214/11-AOAS455

22. Hoerl, A.E., Kennard, R.W.: Ridge regression: Biased estimation for nonorthogonal problems. Technometrics 12(1), 55–67 (Feb 1970). 10.1080/00401706.1970.10488634, http://dx.doi.org/10.1080/00401706.1970.10488634

23. Holland, D., Frei, O., Desikan, R., Fan, C.C., Shadrin, A.A., Smeland, O.B., Sundar, V.S., Thompson, P., Andreassen, O.A., Dale, A.M.: Beyond snp heritability: Polygenicity and discoverability of phenotypes estimated with a univariate gaussian mixture model. PLOS Genetics 16(5), e1008612 (May 2020). 10.1371/journal.pgen.1008612, http://dx.doi.org/10.1371/journal.pgen.1008612

24. Imhof, J.P.: Computing the distribution of quadratic forms in normal variables. Biometrika 48(3/4), 419 (Dec 1961). 10.2307/2332763, http://dx.doi.org/10.2307/2332763

25. Jordan, M.I., Ghahramani, Z., Jaakkola, T.S., Saul, L.K.: An introduction to variational methods for graphical models. Machine Learning 37(2), 183–233 (1999). 10.1023/a:1007665907178, http://dx.doi.org/10.1023/A:1007665907178

26. Kim, S.A., Cho, C.S., Kim, S.R., Bull, S.B., Yoo, Y.J.: A new haplotype block detection method for dense genome sequencing data based on interval graph modeling of clusters of highly correlated snps. Bioinformatics 34(3), 388–397 (Sep 2017). 10.1093/bioinformatics/btx609, http://dx.doi.org/10.1093/bioinformatics/btx609

27. Lehne, B., Lewis, C.M., Schlitt, T.: From snps to genes: Disease association at the gene level. PLoS ONE 6(6), e20133 (Jun 2011). 10.1371/journal.pone.0020133, http://dx.doi.org/10.1371/journal.pone.0020133

28. Liu, H., Tang, Y., Zhang, H.H.: A new chi-square approximation to the distribution of non-negative definite quadratic forms in non-central normal variables. Computational Statistics & Data Analysis 53(4), 853–856 (Feb 2009). 10.1016/j.csda.2008.11.025, http://dx.doi.org/10.1016/j.csda.2008.11.025

29. Logsdon, B.A., Hoffman, G.E., Mezey, J.G.: A variational bayes algorithm for fast and accurate multiple locus genome-wide association analysis. BMC Bioinformatics 11(1) (Jan 2010). 10.1186/1471-2105-11-58, http://dx.doi.org/10.1186/1471-2105-11-58

30. Martin, A.R., Gignoux, C.R., Walters, R.K., Wojcik, G.L., Neale, B.M., Gravel, S., Daly, M.J., Bustamante, C.D., Kenny, E.E.: Human demographic history impacts genetic risk prediction across diverse populations. The American Journal of Human Genetics 107(4), 788–789 (Oct 2020). 10.1016/j.ajhg.2020.08.020, http://dx.doi.org/10.1016/j.ajhg.2020.08.020

31. National Cancer Institute: Genetic simulation resources (gsr) (2024), https://surveillance.cancer.gov/genetic-simulation-resources/

32. Nickl, P.: Bayesian Inference for Regression Models using Nonparametric Infinite Mixtures. Master’s thesis, Technical University of Darmstadt (2020)

33. Pasaniuc, B., Price, A.L.: Dissecting the genetics of complex traits using summary association statistics. Nature Reviews Genetics 18(2), 117–127 (Nov 2016). 10.1038/nrg.2016.142, http://dx.doi.org/10.1038/nrg.2016.142

34. Popejoy, A.B., Fullerton, S.M.: Genomics is failing on diversity. Nature 538(7624), 161–164 (Oct 2016). 10.1038/538161a, http://dx.doi.org/10.1038/538161a

35. Privé, F.: Optimal linkage disequilibrium splitting. Bioinformatics 38(1), 255–256 (Jul 2021). 10.1093/bioinformatics/btab519, http://dx.doi.org/10.1093/bioinformatics/btab519

36. Pruitt, K.D.: Ncbi reference sequence (refseq): a curated non-redundant sequence database of genomes, transcripts and proteins. Nucleic Acids Research 33(Database issue), D501–D504 (Dec 2004). 10.1093/nar/gki025, http://dx.doi.org/10.1093/nar/gki025

37. Schaid, D.J., Chen, W., Larson, N.B.: From genome-wide associations to candidate causal variants by statistical fine-mapping. Nature Reviews Genetics 19(8), 491–504 (May 2018). 10.1038/s41576-018-0016-z, http://dx.doi.org/10.1038/s41576-018-0016-z

38. Sethuraman, J.: A constructive definition of dirichlet priors. Statistica Sinica pp. 639–650 (1994)

39. Sirugo, G., Williams, S.M., Tishkoff, S.A.: The missing diversity in human genetic studies. Cell 177(1), 26–31 (Mar 2019). 10.1016/j.cell.2019.02.048, http://dx.doi.org/10.1016/j.cell.2019.02.048

40. Sollis, E., Mosaku, A., Abid, A., Buniello, A., Cerezo, M., Gil, L., Groza, T., Güneş, O., Hall, P., Hayhurst, J., Ibrahim, A., Ji, Y., John, S., Lewis, E., MacArthur, J.A.L., McMahon, A., Osumi-Sutherland, D., Panoutsopoulou, K., Pendlington, Z., Ramachandran, S., Stefancsik, R., Stewart, J., Whetzel, P., Wilson, R., Hindorff, L., Cunningham, F., Lambert, S.A., Inouye, M., Parkinson, H., Harris, L.W.: The NHGRI-EBI GWAS catalog: knowledgebase and deposition resource. Nucleic Acids Research 51(D1), D977–D985 (Nov 2022). 10.1093/nar/gkac1010, http://dx.doi.org/10.1093/nar/gkac1010

41. Solovieff, N., Cotsapas, C., Lee, P.H., Purcell, S.M., Smoller, J.W.: Pleiotropy in complex traits: challenges and strategies. Nature Reviews Genetics 14(7), 483–495 (Jun 2013). 10.1038/nrg3461, http://dx.doi.org/10.1038/nrg3461

42. Spain, S.L., Barrett, J.C.: Strategies for fine-mapping complex traits. Human Molecular Genetics 24(R1), R111–R119 (Jul 2015). 10.1093/hmg/ddv260, http://dx.doi.org/10.1093/hmg/ddv260

43. Stearns, F.W.: One hundred years of pleiotropy: A retrospective. Genetics 186(3), 767–773 (Nov 2010). 10.1534/genetics.110.122549, http://dx.doi.org/10.1534/genetics.110.122549

44. Stephens, M.: False discovery rates: a new deal. Biostatistics p. kxw041 (Oct 2016). 10.1093/biostatistics/kxw041, http://dx.doi.org/10.1093/biostatistics/kxw041

45. Sudlow, C., Gallacher, J., Allen, N., Beral, V., Burton, P., Danesh, J., Downey, P., Elliott, P., Green, J., Landray, M., Liu, B., Matthews, P., Ong, G., Pell, J., Silman, A., Young, A., Sprosen, T., Peakman, T., Collins, R.: Uk biobank: An open access resource for identifying the causes of a wide range of complex diseases of middle and old age. PLOS Medicine 12(3), e1001779 (Mar 2015). 10.1371/journal.pmed.1001779, http://dx.doi.org/10.1371/journal.pmed.1001779

46. Svishcheva, G.R., Belonogova, N.M., Zorkoltseva, I.V., Kirichenko, A.V., Axenovich, T.I.: Gene-based association tests using gwas summary statistics. Bioinformatics 35(19), 3701–3708 (Mar 2019). 10.1093/bioinformatics/btz172, http://dx.doi.org/10.1093/bioinformatics/btz172

47. Thompson, W.K., Wang, Y., Schork, A.J., Witoelar, A., Zuber, V., Xu, S., Werge, T., Holland, D., Andreassen, O.A., Dale, A.M.: An empirical bayes mixture model for effect size distributions in genome-wide association studies. PLOS Genetics 11(12), e1005717 (Dec 2015). 10.1371/journal.pgen.1005717, http://dx.doi.org/10.1371/journal.pgen.1005717

48. Tibshirani, R.: Regression shrinkage and selection via the lasso. Journal of the Royal Statistical Society Series B: Statistical Methodology 58(1), 267–288 (Jan 1996). 10.1111/j.2517-6161.1996.tb02080.x, http://dx.doi.org/10.1111/j.2517-6161.1996.tb02080.x

49. Turley, P., Walters, R.K., Maghzian, O., Okbay, A., Lee, J.J., Fontana, M.A., Nguyen-Viet, T.A., Wedow, R., Zacher, M., Furlotte, N.A., Magnusson, P., Oskarsson, S., Johannesson, M., Visscher, P.M., Laibson, D., Cesarini, D., Neale, B.M., Benjamin, D.J.: Multi-trait analysis of genome-wide association summary statistics using mtag. Nature Genetics 50(2), 229–237 (Jan 2018). 10.1038/s41588-017-0009-4, http://dx.doi.org/10.1038/s41588-017-0009-4

50. Uffelmann, E., Huang, Q.Q., Munung, N.S., de Vries, J., Okada, Y., Martin, A.R., Martin, H.C., Lappalainen, T., Posthuma, D.: Genome-wide association studies. Nature Reviews Methods Primers 1(1) (Aug 2021). 10.1038/s43586-021-00056-9, http://dx.doi.org/10.1038/s43586-021-00056-9

51. Urbut, S.M., Wang, G., Carbonetto, P., Stephens, M.: Flexible statistical methods for estimating and testing effects in genomic studies with multiple conditions. Nature Genetics 51(1), 187–195 (Nov 2018). 10.1038/s41588-018-0268-8, http://dx.doi.org/10.1038/s41588-018-0268-8

52. Visscher, P.M., Wray, N.R., Zhang, Q., Sklar, P., McCarthy, M.I., Brown, M.A., Yang, J.: 10 years of gwas discovery: Biology, function, and translation. The American Journal of Human Genetics 101(1), 5–22 (Jul 2017). 10.1016/j.ajhg.2017.06.005, http://dx.doi.org/10.1016/j.ajhg.2017.06.005

53. Visscher, P.M., Yengo, L., Cox, N.J., Wray, N.R.: Discovery and implications of polygenicity of common diseases. Science 373(6562), 1468–1473 (Sep 2021). 10.1126/science.abi8206, http://dx.doi.org/10.1126/science.abi8206

54. Wagner, G.P., Zhang, J.: The pleiotropic structure of the genotype–phenotype map: the evolvability of complex organisms. Nature Reviews Genetics 12(3), 204–213 (Feb 2011). 10.1038/nrg2949, http://dx.doi.org/10.1038/nrg2949

55. Wainwright, M.J., Jordan, M.I.: Graphical models, exponential families, and variational inference. Foundations and Trends® in Machine Learning 1(1–2), 1–305 (2007). 10.1561/2200000001, http://dx.doi.org/10.1561/2200000001

56. Wang, G., Sarkar, A., Carbonetto, P., Stephens, M.: A simple new approach to variable selection in regression, with application to genetic fine mapping. Journal of the Royal Statistical Society Series B: Statistical Methodology 82(5), 1273–1300 (Jul 2020). 10.1111/rssb.12388, http://dx.doi.org/10.1111/rssb.12388

57. Wu, T.T., Chen, Y.F., Hastie, T., Sobel, E., Lange, K.: Genome-wide association analysis by lasso penalized logistic regression. Bioinformatics 25(6), 714–721 (Jan 2009). 10.1093/bioinformatics/btp041, http://dx.doi.org/10.1093/bioinformatics/btp041

58. Yang, J., Ferreira, T., Morris, A.P., Medland, S.E., Madden, P.A.F., Heath, A.C., Martin, N.G., Montgomery, G.W., Weedon, M.N., Loos, R.J., Frayling, T.M., McCarthy, M.I., Hirschhorn, J.N., Goddard, M.E., Visscher, P.M.: Conditional and joint multiple-snp analysis of gwas summary statistics identifies additional variants influencing complex traits. Nature Genetics 44(4), 369–375 (Mar 2012). 10.1038/ng.2213, http://dx.doi.org/10.1038/ng.2213

59. Zhang, Y., Qi, G., Park, J.H., Chatterjee, N.: Estimation of complex effect-size distributions using summary-level statistics from genome-wide association studies across 32 complex traits. Nature Genetics 50(9), 1318–1326 (Aug 2018). 10.1038/s41588-018-0193-x, http://dx.doi.org/10.1038/s41588-018-0193-x

60. Zhao, Y., Zhu, H., Lu, Z., Knickmeyer, R.C., Zou, F.: Structured genome-wide association studies with bayesian hierarchical variable selection. Genetics 212(2), 397–415 (Apr 2019). 10.1534/genetics.119.301906, http://dx.doi.org/10.1534/genetics.119.301906

61. Zhou, X., Carbonetto, P., Stephens, M.: Polygenic modeling with bayesian sparse linear mixed models. PLoS Genetics 9(2), e1003264 (Feb 2013). 10.1371/journal.pgen.1003264, http://dx.doi.org/10.1371/journal.pgen.1003264

62. Zhu, X., Stephens, M.: Bayesian large-scale multiple regression with summary statistics from genome-wide association studies. The Annals of Applied Statistics 11(3) (Sep 2017). 10.1214/17-aoas1046, http://dx.doi.org/10.1214/17-AOAS1046

63. Zou, H., Hastie, T.: Regularization and variable selection via the elastic net. Journal of the Royal Statistical Society Series B: Statistical Methodology 67(2), 301–320 (Mar 2005). 10.1111/j.1467-9868. 2005.00503.x, http://dx.doi.org/10.1111/j.1467-9868.2005.00503.x

64. Zou, Y., Carbonetto, P., Wang, G., Stephens, M.: Fine-mapping from summary data with the “sum of single effects” model. PLOS Genetics 18(7), e1010299 (Jul 2022). 10.1371/journal.pgen.1010299, http://dx.doi.org/10.1371/journal.pgen.1010299

