## Appendices for "ML-MAGES: A machine learning framework for multivariate genetic association analyses with genes and effect size shrinkage"

### A Appendices

#### A.1 UK Biobank data

The database of UK Biobank contains the whole genome sequencing for about 500,000 participants, with genotyping data from 800,000 genome-wide variants and imputation to 90 million variants [45]. Our study is conducted under an approved project from UK Biobank. We restrict our analysis to genotyped variants of 22 autosomes from individuals of European ancestry, and go through a series of quality control (QC) on the variants and samples. `PLINK2` [12] is used to perform all QC steps as well as GWA analyses. We use two quantitative traits from UK Biobank in our study, the direct low-density lipoproteins cholesterol (LDL, Data-Field 30780) and the high-density lipoprotein cholesterol (HDL, Data-Field 30760).

*Quality control of the genetic data.* For QC, we filter out variants with minor allele frequency below 0.01, missing call rates exceeding 0.01, Hardy-Weinberg equilibrium exact test p-value below 1e-6, and with one or more multi-character allele codes or single-character allele codes outside of 'ACGT' (`PLINK2` command `-maf 0.01 -hwe 1e-6 -geno 0.01 -snps-only just-acgt`). We also exclude ambiguous SNPs, i.e., those with complementary alleles, either 'C/G' or 'A/T' SNPs.

We exclude samples with missing call rates exceeding 0.05 (`-mind 0.05`), as well as samples with putative aneuploidy, samples with excess third-degree relatives, outliers based on heterozygosity and missing rates, and samples that withdrawal from the study. We restrict our analysis to samples that have a reported genetic ethnic grouping of "White British" from the UK Biobank database (Data-Field 21000) that corresponds to the commonly labeled "European ancestry".

After QC, a total of 489,953 variants remained for a total of 334,851 people (155402 males, 179449 females). The total genotyping rate is 0.954831. In calculations that involve using the genotype matrix, any remaining missing genotype is imputed by the average value of the non-missing ones.

*GWA and LD calculation.* In GWA studies, we include sex, age, body mass index (BMI), and the top 20 PCs from principal component analysis on the genotype data representing population structure as the covariates (a total of 23 covariates). PCA is performed using `flashpca` [1] on the variants pruned using `PLINK2` command `-indep-pairwise 100 10 0.1`. In-sample LD is calculated on UKB data using command `-r2 square`.

#### A.2 Input to *ML-MAGES*: GWA summary statistics and LD

Suppose the individual-level GWA data contains genotypes of $m$ variants (SNPs) from a set of $n$ diploid individuals, where each sample contains two sets of chromosomes. SNP genotypes are typically coded as 0, 1 or 2 to represent the number of non-reference alleles at each position. Let the genotype matrix be $X : n \times m$, where $X_{ij} \in \{0, 1, 2\}$, and let $x_j$ be the $j$-th column of the matrix denoting the genotypes of variant $j$. Let $y \in \mathbb{R}^n$ be the phenotype, and assume it is standardized. In this study, we only consider continuous traits.

Let $\beta \in \mathbb{R}^m$ denote some unknown true effect sizes of the variants. Under the assumption in which effect of each copy of an allele on the trait is assumed to be additive (the "additive model" [11,50]),

$$y = X\beta + \epsilon, \tag{4}$$

where $\epsilon \in \mathbb{R}^n$ is noise following $\epsilon_i \sim \mathcal{N}(0, \eta^2)$.

In GWA studies, the approach is to perform simple linear regression $y = x_j\beta_j + \epsilon$ on each genetic variant across the genome on $y$, the trait of interest. The estimated effect is denoted as $\hat{\beta}_j$, with standard error $s_j$. The regression commonly includes covariates like age, sex, and top principal components of the genotype matrix which incorporate the population structure. For simplicity disregarding the covariates, an observed effect $\hat{\beta}_j$ from GWA studies can be estimated via ordinary least squares as

$$\hat{\beta}_j = (x_j^T x_j)^{-1} x_j^T y. \tag{5}$$

Assuming that $X$ is column-standardized, the $m \times m$ in-sample LD matrix, or the genetic correlation matrix, can be obtained as $R = \frac{1}{n} X^T X$. Each entry $R_{jj'}$ is the linkage disequilibrium (LD) between a pair of SNPs $j$ and $j^{'}$, calculated as the squared correlation based on genotypic allele counts. The LD score of a SNP $j$, which quantifies the amount of genetic variation tagged by the variant, is the sum of its squared correlations with all other SNPs and can be calculated as $\ell_j = \sum_{j'=1, j' \neq j}^{m} R_{jj'}$ [9].

Due to privacy concerns, the individual-level genotype data $X$ is often not publicly accessible. However, there are abundant summary-level data of GWA summary statistics in public database like GWAS catalog [10,40]. Methods only requiring summary-level data are therefore preferable to ensure their general applicability [33]. Similar to many existing methods [47,44,62,59], we intend our method to use only summary-level data and LD information as input. LD may be obtained on out-of-sample reference population of the same ancestry (e.g., from 1000 Genomes Project [20]) when individual-level data is not available.

##### A.3 Related methods for effect size shrinkage

*Shrinkage via mixture models.* A group of methods use Gaussian mixture models to introduce sparsity (i.e., zeros) into effect sizes, which is essentially shrinking the inflated effects [29,21,61,47,44,62,59,23]. Many methods use a mixture of two components, usually zero-mean Gaussian distributions, with one of them having a point mass at zero to model the nearly zero effects from null SNPs [29,21,61,47,62,59,23]. [59] also extends this approach to three mixture components, allowing some non-null variants to have distinctly larger effects. The adaptive shrinkage method *ash* by [44] uses a mixture of $K$ components for the true effect size distribution with a point mass for truly null SNPs. [51] introduces the multivariate adaptive shrinkage method *mash* which uses multivariate Gaussian components, where the covariance matrices of the mixture components are designed to captures patterns of multiple groups of effects. In both *ash* and *mash*, the variances or the covariance matrices of the $K$ mixture components are generated and fixed for a pre-specified number $K$, while the mixture weights are estimated to show the distributions of effects. Both methods work with summary-level data and do not explicitly account for LD.

*Fine-mapping.* The goal of fine-mapping is similar to the goal of shrinkage, but on a much smaller scale: to pinpoint, within some small trait-associated region from a GWA study, a few variants that very likely contain a causal one [42,33,37]. The focal regions are usually small chromosomal segments that show a lots of significant hits in GWA, where the aggregation of hits is likely due to high LD between the SNPs. Statistical methods for fine-mapping include identifying the lead variant in the associated region based on P-values or LD and Bayesian approaches which assign posterior probabilities of causality to each variant [42]. *SuSiE* [56] formulates the task as a variable selection problem and uses a Bayesian model for sparse multiple regression to identify variables (variants in fine-mapping) with non-zero effect; its extension *SuSiE-RSS* [64] allows the use of only summary data as input. However, in fine-mapping, the true effects are highly sparse; fine-mapping techniques rarely scale to the genome-wide shrinkage task, especially if the trait is highly polygenic. Additionally, Bayesian inference, often performed through MCMC, can become computationally prohibitive genome-wide.

*Shrinkage as regularized regression: the gene-$\epsilon$ approach.* The per-SNP observed GWA effects, $\hat{\beta}$, are sometimes called marginal effects, and the true effects in the additive model, $\beta$, are sometimes called joint effects. A simple relationship between marginal and joint effects has been derived and used in many studies [58,62,59,23]:

$$\mathbb{E}[\hat{\beta}_j] = \sum_{j'=1}^{m} R_{jj'} \beta_{j'}, \tag{6}$$

or in matrix form, $\mathbb{E}[\hat{\beta}] = R\beta$. Additional scaling factors are sometimes included for each variant in the linear summation according to their SNP heterozygosity [23] or GWA standard errors [62]. The observed effect of a variant is a weighted sum of true effects of all variants, weighted by their genetic correlations (captured in $R_{jj'}$) to the focal variant.

Based on this linear relationship, *gene-$\epsilon$* [14] performs shrinkage through a regularized regression, comparing LASSO [48], ridge regression [22], and elastic net [63]. Elastic net is shown to work best. The regularized $\tilde{\beta}$ is obtained by

$$\arg\min_{\tilde{\beta}} \frac{1}{m} \|\hat{\beta} - R\tilde{\beta}\|_2^2 + \lambda(\tilde{\beta}), \tag{7}$$

where $\|\cdot\|_1$ is the L1-norm, $\|\cdot\|_2$ is the L2-norm, and $\lambda(\cdot)$ is a regularization term that penalizes for the non-zero values in $\tilde{\beta}$. The elastic net regularization has

$$\lambda(\tilde{\beta}) = \lambda_1 \left(\lambda_2 \|\tilde{\beta}\|_1 + (1-\lambda_2)\|\tilde{\beta}\|_2^2/2\right) \tag{8}$$

where $\lambda_1$ is a multiplier controlling the overall strength of the penalty, and $\lambda_2$ is the mixing parameter controlling the combination of L1 and L2. $\lambda_2 = 1$ corresponds to the LASSO regularization and $\lambda_2 = 0$ corresponds to ridge regression.

##### A.4 Synthetic data generation and training data preparation

Generating realistic genome-wide genotype simulations is a complex task, as it involves producing both realistic allele frequency distributions and LD patterns. Simulating genotypes from scratch using a genetic data simulator (see [31] for a list of resources) is beyond the scope of this study. Simulation from genotype data is a common practice in genetic studies [62,59,56,14,64]. Here, we first sub-sample genotypes from real chromosomal data, and then use simulations to generate synthetic true effect sizes and trait values following the additive model in Eq. 4.The observed effects are obtained using simple regressions on the synthetic trait. We vary parameters in the simulations such as the narrow-sense heritability, which is the ratio of additive genetic variance to the total phenotypic variance, and the total number of associated variants. The real chromosomal data used in simulation come from UK Biobank.

We take two approaches to simulate the true effects, one based on just SNP-level data, and one based on gene-level information as well. We refer to these as "SNP-only simulation" and "gene-level simulation", respectively. For the SNP-only simulation, we subsample regions of variants in a chromosome from a subset of individuals to get genotype matrices. We then simulate the true effects and noise terms, and obtain simulated traits while fixing the narrow-sense heritability. For the gene-level simulation, instead of sampling random regions from the chromosomes, an entire chromosome—randomly chosen—is used to simulate the data. Two traits are simulated evaluate our multi-trait analysis. From all genes in the chromosome, some are chosen to be truly associated to one of the two traits, and some are chosen to be associated to both. A fraction of variants in those chosen genes are set to be associated, i.e., with non-zero true effects. Simulation steps are detailed below.

Next, GWA studies are performed on the simulated traits to generate simulated univariate summary-level data. By simulation, we generate both the ground truth effects, $\beta^{(s)}$, and the GWA summary statistics, $\hat{\beta}^{(s)}$ and $s^{(s)}$. SNP-only simulation data is then used as the training and validation data for training the NNs. We also use NP-only simulations to benchmark the performance of different methods for shrinkage. We use different chromosomes for simulating training data and validation data to ensure they do not overlap. Gene-level simulation is not used for training. We use it solely for performance comparison end-to-end, including shrinkage, clustering, and gene-level analysis (Fig. 2).

An issue we encountered is that simulated observed effects do not closely follow the distribution of real GWA effects. Despite the widely used additive model (Eq. 4), the true generative processes underling our genome is much more complicated than those suggested by the theoretical models. However, supervised machine learning models rely on a good alignment between training and test data to perform well. We bridge this gap between simulated data and real data by applying transformations on the simulated values, both $\hat{\beta}^{(s)}$ and $s^{(s)}$.

We manipulate the simulated GWA summary statistics to better match the distribution of the real ones for a good alignment between training and test data. We do this by applying transformations on the simulated $\hat{\beta}^{(s)}$ and $s^{(s)}$. We notice that the distribution of real GWA effects $\hat{\beta}$ follows a Laplace distribution more closely than a normal one, so we fit a zero-centered Laplace distribution on $\hat{\beta}$ of the traits of interest. We then transform simulated $\hat{\beta}^{(s)}$ values so that the empirical cumulative distribution follows this fitted Laplace distribution. For the standard errors, we simply re-scale them so that the scales of synthetic $s^{(s)}$ match the real ones. In this way, the simulated values, after transformation, distribute similarly to the real ones, and we are ready to use them for model training.

*SNP-only simulation.* We subsample several regions of $m^{(s)} = 1000$ variants in a chromosome from a subset of $N^{(s)} = 10000$ individuals. Each sampling gives a genotype data matrix $X^{(s)}$ of size $m^{(s)} \times N^{(s)}$. The SNP-only simulation procedure for each sampled $X^{(s)}$ is as follows:

1. Set the heritability of the variants $h^2$ to be uniformly from $[0.1, 0.9]$.
2. Randomly pick a set of causal SNPs, $C$, with the number of causal variants $|C|$ sampled as integers uniformly from $[1, 0.05 * m^{(s)}] = [1, 50]$.
3. Simulate true effects as $\beta_j^{(s)} \sim \mathcal{N}(0, 1)$ for $j \in C$ and $\beta_j^{(s)} = 0$ otherwise.
4. Re-scale the simulated effects $\beta^{(s)}$ so that $\mathrm{Var}(X^{(s)}\beta^{(s)}) = h^2$.
5. Simulate a noise vector $\epsilon^{(s)}$ of size $N^{(s)}$, with $\epsilon_j^{(s)} \sim \mathcal{N}(0, 1 - h^2)$.
6. Simulate synthetic traits as $y^{(s)} = X^{(s)}\beta^{(s)} + \epsilon^{(s)}$.
7. Perform simple regressions on $m^{(s)}$ variants separately:
$$y^{(s)} = x_j^{(s)}\beta_j^{(s)} + \epsilon^{(s)},$$
giving an observed effect $\hat{\beta}_j^{(s)}$ and its corresponding standard error $s_j^{(s)}$ for each variant.
8. Take the LD of the simulated data, $R^{(s)}$, as the $m^{(s)} \times m^{(s)}$ matrix block that corresponds to the subset of sampled variants from the full LD matrix of the chromosome calculated using the real genotype data.
9. Transform the simulated $\hat{\beta}^{(s)}$ and rescale $s^{(s)}$ to distribute similarly to the real ones.
10. Construct features $\Omega^{(s)}$ from $\hat{\beta}^{(s)}$, $s^{(s)}$, and $R^{(s)}$ for supervised learning following Eq. 2.

The transformation of the simulated betas $\hat{\beta}^{(s)}$ are done by first fitting a Laplace distribution to the real GWA effects (here we use those of LDL and HDL), and then transforming the simulated data's empirical distribution to match that of the fitted Laplace. The scaling of the simulated standard errors $s^{(s)}$ follows

$$\frac{s^{(s)} - a^{(s)}}{b^{(s)} - a^{(s)}}(b - a) + a, \tag{9}$$

where $(a, b)$, and $(a^{(s)}, b^{(s)})$ are the (1%,99%)-quantile values of the real $s$ and the simulated $s^{(s)}$. The quantile values instead of the minimum and maximum are used to make the scaling more robust to outliers.

For the training data, we run 200 simulations each using chromosomes 18, 19, 21, and 22, resulting in a total of $200 \times 4 \times 1000 = 800,000$ data points. For the validation data, we run 200 simulations using chromosome 20, resulting in a total of $200 \times 1000 = 200,000$ data points. The validation data is also used in performance comparisons.

We also separately generate a training set comprised of 100 simulations each based on chromosomes 1, 7, 13, and 19, and a validation set comprised of 100 simulations each based on chromosomes 3 and 14. The training data has size 400,000 and the validation data has size 200,000. The same neural-network models are also trained and validated on these data, and they are assessed on the previous simulation data, showing similar performances. Results are not shown.

*Gene-level simulation.* Gene-level simulation is based on 15,250 genotyped variants on chromosome 15 from UK Biobank. There are 433 genes with a number of SNPs between 2 and 265. We generate the causality patterns of two simulated traits. For simulation, we randomly sample 11 to 13 out of the 274 genes (approximately 5%) with at least 6 SNPs to be truly associated to either one or both of the traits. Either 30% of all the SNPs in each causal gene or two SNPs, whichever comes larger, are chosen to be causal. For those that are associated to both, true effects are drawn from a zero-mean bivariate normal that result in a high correlation—either positive or negative—between them. For variants that are only associated to one trait, we draw the true effects from a zero-mean univariate normal. The narrow-sense heritability of both traits are fixed at $h^2 = 0.8$. The rest of the simulation procedure follows the same steps 4 to 10 as in the SNP-only simulation. We generate 100 gene-level simulations, each containing the effects of 15,250 variants for two traits with some shared associated genes, where a total number of associated SNPs for either trait varies between 50 and 150. Altogether there are 200 single-trait simulations, and they are used for performance comparisons of shrinkage methods (Fig. 2).

##### A.5 LD block decomposition to reduce the problem size

Both elastic net [14] and *SuSiE-RSS* [56,64] are computationally expensive when shrinking many variants, e.g., all genotyped variants from an entire chromosome—even for the shortest autosome. Performing shrinkage using these methods on the entire LD matrix of a chromosome is inefficient.

Variants far apart on the chromosome have relatively low LD, making the LD matrix close to a block-diagonal matrix. It is therefore common to split the genome into nearly independent blocks of LD. While operations on the full LD can be computationally expensive, applying them to smaller matrices can make these operations faster and parallelizable [5,26,35]. We decompose the LD matrix of each chromosome into multiple LD blocks and perform shrinkage on each LD block separately. This allows us both to conduct performance comparison and to reduce the feature space of NN models. The results obtained on each LD block are then aggregated together.

We adopt a method from [5] to get approximately independent LD blocks. Each antidiagonal term of the LD is represented by the sum of its elements. A low sum value indicates that variants on two sides of the antidiagonal are weakly linked. Next, a low-pass filter is passed on the antidiagonal sums to reduce noise and a desired number of minima are chosen to be the candidate segment boundaries, followed by a local search in the proximity of each minimum for fine-tuning. The number of blocks is chosen so that the each block contains around 1000 variants.

##### A.6 Neural network architectures and training settings

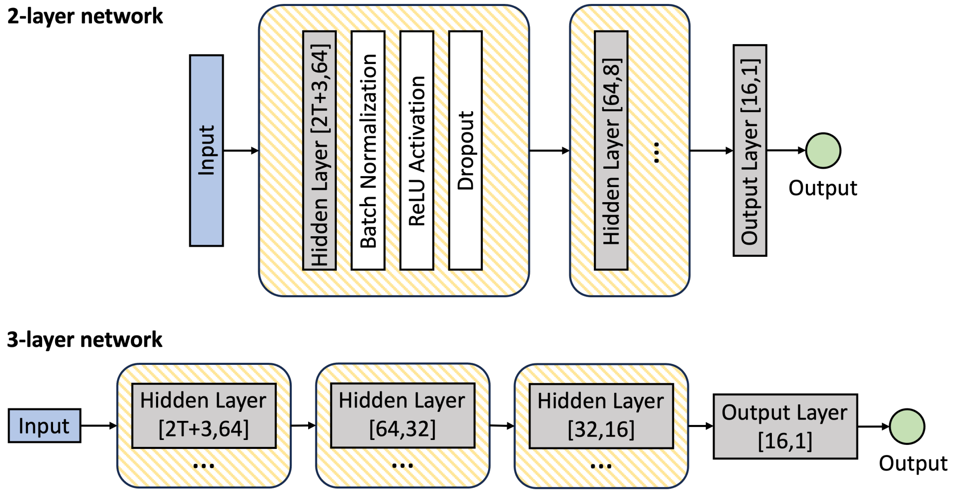


**Fig. 5. Architectures of the neural network models used for shrinkage.** Input consists of observed effect $\hat{\beta}_i$, standard error $s_i$, LD score $\ell_i$ of the focal variant $i$, as well as observed effects $\hat{\beta}_j$ and LD values $r_{ij}$ of the top $T$ variants that are in highest LD ($r_{ij}$) with the focal variant $i$. Output is the regularized effect $\tilde{\beta}_i$, aiming at approximate the unknown true effect $\beta i$. Input and output dimensions of each layer are included in brackets. The dropout layer is not included in the last hidden layer in each model.

In our network models, each fully-connected hidden layer is followed by a batch normalization layer and a ReLU activation function. The final layer, which is also a fully-connected linear layer, outputs the desired regularized effect. To prevent over-fitting, drop-out layers with drop out rate of 0.2 are also included. A graphical representation of the architectures are shown in Fig. A.6.

For training, we use the Adam optimizer with a learning rate of $10^{-4}$ and MSE loss for the objective. We train each model with a batch size of 100 for a maximum of 500 epochs and implement early-stopping to prevent over-fitting. The training always terminates early.

Neural-network based shrinkage performs consistently across network architectures with different number of layers and input layer sizes, and no specific architecture excels at all tasks. In this study, we compare a few architectures and simply choose the 2-layer and 3-layer architectures using top 15 variants for features as demonstrations. We do not delve into the exploitation of model parameters and training hyperparameters, as our focus lies primarily on demonstrating the practical application of such deep supervised learning methods for shrinkage. In general, while more complex models possess greater computational power, they are also subject to a higher computational burden and an increased risk of overfitting to the specific training data. Therefore, we recommend selecting an architecture that maintains a good balance between complexity and generalization, ideally validated on small subsets of data.

##### A.7 Probability density functions of distributions

Below are the probability density functions of the distributions used in the inference of the zero-mean infinite-mixture model. A categorical distribution parameterized by $\pi$ of size $K$ is denoted as $\mathrm{Cat}(\pi)$, with pdf

$$f(z|\pi) = \mathrm{Cat}(z|\pi) = \prod_{k=1}^{K} [\pi_k]^{z_k}. \tag{10}$$

A Beta distribution with shape parameters $a$ and $b$ is denoted as $\mathrm{Beta}(a, b)$, with pdf

$$f(v|a,b) = \mathrm{Beta}(v|a,b) = \frac{1}{B(a,b)} v^{a-1}(1-v)^{b-1}, \tag{11}$$

where $B(a,b) = \frac{\Gamma(a)\Gamma(b)}{\Gamma(a+b)}$ is the beta function and $\Gamma(\cdot)$ is the gamma function.

A Wishart distribution with $\nu$ degrees of freedom and a $p \times p$ symmetric positive-definite scale matrix $L$ is denoted as $\mathcal{W}(L, \nu)$, with pdf

$$f(A|L,\nu) = \mathcal{W}(A|L,\nu) = W(L,\nu)|A|^{\frac{\nu-p-1}{2}} \exp\{-\frac{1}{2}\mathrm{Tr}(L^{-1}A)\}, \tag{12}$$

where $W(L,\nu) = \frac{1}{2^{\nu p/2}|L|^{\nu/2}\Gamma_p(\nu/2)} = 2^{-\nu p/2}\,|L|^{-\nu/2}\left(\pi^{\frac{p(p-1)}{4}}\prod_{j=1}^{p}\Gamma(\frac{\nu+1-i}{2})\right)^{-1}$.

##### A.8 Variational inference

Suppose there are $p$ traits and $J \leq m$ non-zero effects—non-zero for any of the trait—after shrinkage. We want to cluster the $J$ variants according to their associations represented by the effect sizes. The known data is $\{\gamma_i\}_{i=1}^{J}$, where

$$\{\gamma_i\} = \{\tilde{\beta}_j \in \mathbb{R}^p | \tilde{\beta}_j \not\approx \mathbf{0} \text{ for } j = 1, \ldots, m\}. \tag{13}$$

These are input to the clustering. The latent indicator variables $\{z_i\}_{i=1}^{J}$ are what we want the clustering algorithm to output, where

$$z_{ik} = \begin{cases} 1 \text{ if variant } i \text{ belongs to cluster } k \\ 0 \text{ otherwise} \end{cases}, \tag{14}$$

for a total of $K$ clusters to be determined by the model.

Under a zero-mean infinite-mixture model,

$$\gamma_i \sim \sum_{k=1}^{\infty} \pi_k \mathcal{N}(0, A_k^{-1}) \tag{15}$$

$$z_i \sim \mathrm{Cat}(\pi), \tag{16}$$

$$\pi_k = v_k \prod_{\ell=1}^{k-1}(1 - v_\ell). \tag{17}$$

Note that $v$ deterministically gives the values of $\pi$. We use the priors

$$v_k \sim \mathrm{Beta}(a_0, b_0) = \mathrm{Beta}(1, \alpha) \tag{18}$$

and

$$A_k \sim \mathcal{W}(L_0, \nu_0), \tag{19}$$

for $v$ and $A$, where $\text{Beta}(a, b)$ denotes a Beta distribution and $\mathcal{W}(L, \nu)$ denotes a Wishart distribution, and $a_0, b_0, L_0, \nu_0$ are the hyperparameters.

We need to approximate the intractable posterior $p(z, v, A|\gamma)$ of our model. Variational inference (VI) [25,55] is a technique used in Bayesian statistics and machine learning to approximate complex probability distributions by a "closest" simpler variational distribution, typically from a predefined family of simpler distributions, which is parameterized by variational parameters.

We approximate the posterior distribution of our model using VI. The variational variables are $\{z, v, A\}$—the latent indicators $z$, the parameters for mixture weights $v$, and the precision matrices of the Gaussiance distributions $A$—and the observed data are the non-zero regularized effects $\{\gamma\}$. A graphical representation of the model is included in Fig. A.8. Specifically, we use the technique of mean-field inference and coordinate-ascent optimization to perform inference. Under mean-field assumptions, the variational distribution factorize over partitions of mutually independent latent variables. Coordinate ascent variational inference (CAVI) is a typical method to optimize the evidence lower bound (ELBO), in which the variational approximation of each partition of the latent variables is optimized in turn while holding the others fixed. This is sometimes also called variational expectation maximization, where coordinate updates on local variables correspond to the E-step and the updates on global variables correspond to the M-step in the classic EM. Optimization steps are detailed below.

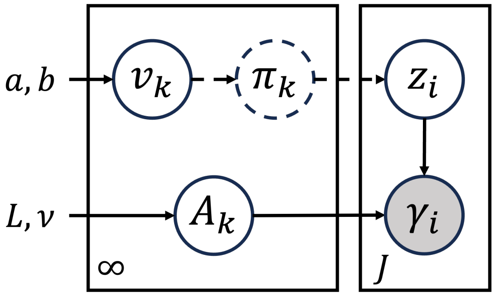


**Fig. 6.** A graphical representation of the zero-mean infinite-mixture (Eq. 3) using plate notation. Nodes are random variables and edges represent the dependence between them. The observed variable is shaded. Uncircled letters are hyperparameters. Dashed lines indicate that $z$ is dependent on $v$ through $\pi$, but $\pi$ is not explicitly included as a variational variable in the inference since it can be deterministically represented by $v$.

*VI for the zero-mean infinite Gaussian mixture model.* The likelihood of data given the model is

$$p(\gamma|z, A) = \prod_{i=1}^{J} \prod_{k=1}^{\infty} \left[\mathcal{N}(0, A_k^{-1})\right]^{z_{ik}} . \tag{20}$$

The conjugate prior on variational variables is

$$\begin{aligned} p(z, \pi, A) &= p(z|\pi)\, p(v)\, p(A) \\ &= \prod_{i=1}^{J} \text{Cat}(z_i|\pi) \prod_{k=1}^{\infty} \text{Beta}(v_k|a_0, b_0) \prod_{k=1}^{\infty} \mathcal{W}(A_k|L_0, \nu_0) \\ &= \prod_{i=1}^{J} \prod_{k=1}^{\infty} \left[ v_k \prod_{\ell=1}^{k-1} (1 - v_\ell) \right]^{z_{ik}} \times \prod_{k=1}^{\infty} \frac{v_k^{a_0-1}(1-v_k)^{b_0-1}}{B(a_0, b_0)} \\ &\quad \times \prod_{k=1}^{\infty} W(L_0, \nu_0) |A_k|^{\frac{\nu_0 - p - 1}{2}} \exp\{-\frac{1}{2}\text{Tr}(L_0^{-1} A_k)\}. \end{aligned} \tag{21}$$

The target distribution is the posterior $p(z, v, A|x)$, and the variational distribution is $q(z, v, A)$. We use mean-field approximation and coordinate-ascent optimization to perform the inference. That is, we assume that $q$ can factorize over parameters $\theta = \{z, v, A\}$. The mean field approximation of the posterior $p(z, v, A|\gamma)$ is

$$\begin{aligned} p(z,v,A|\gamma) \approx q(z,v,A) &= q(z)q(v)q(A) \\ &= \prod_{i=1}^{J} \mathrm{Cat}(z_i|r_i) \prod_{k=1}^{K-1} \mathrm{Beta}(v_k|a_k,b_k) \prod_{k=1}^{K} \mathcal{W}(A_k|L_v,\nu_k). \end{aligned} \tag{22}$$

In the approximation distribution, $K$ is set to an appropriately large value, e.g., $K = 20$. The variational posterior truncated at this upper limit $K$ can be a reasonable approximation to the infinite mixture [6].

In coordinate-ascent variational inference (CAVI), we iterate over each of the variables $z$, $v$, and $A$. For each variable set $t$, we set

$$q^*(t) \propto \exp \mathbb{E}_{q_{\theta_{\neq t}}} \left[\log p(t|\theta_{\neq t}, \mathbf{b})\right], \tag{23}$$

while keep all other $\{q_s(\cdot)\}_{s\neq t}$ fixed, where $\theta_{\neq t}$ denotes the sets of variables excluding $t$. The iteration stops when the algorithm converges to a local optimum of the non-convex ELBO objective. This can be thought of as a variational EM algorithms, where the variational E-step involves updating $q(z)$, and the variational M-step involves updating $q(v)$ and $q(A)$. The derivation of the optimization steps is similar to that of a non-zero multivariate mixture in [32].

*Variational E-step.* Following CAVI, in each iteration, $q(z)$ is updated as

$$\begin{aligned} \log q(z) &= \mathbb{E}_{q(v,A)}[\log p(\gamma, z, v, A)] + C \\ &= \mathbb{E}\left[\log \prod_{i=1}^{J}\prod_{k=1}^{\infty} \left[\mathcal{N}(\gamma_i|0, A_k^{-1})\right]^{z_{ik}}\right] + \mathbb{E}\left[\log \prod_{i=1}^{J} \mathrm{Cat}(z_i|\pi)\right] + C \\ &= \sum_{i=1}^{J}\sum_{k=1}^{K} z_{ik} \log \rho_{ik} + C. \end{aligned} \tag{24}$$

where $\mathbb{E}[\cdot]$ is with respect to $q(v, A)$, $C$ denotes some constant, and

$$\log \rho_{ik} = \frac{1}{2}\mathbb{E}\left[\log|A_k|\right] - \frac{p}{2}\log(2\pi) - \frac{1}{2}\mathbb{E}[\gamma_i^T A_k \gamma_i] + \mathbb{E}[\log v_k] + \sum_{j=1}^{k-1} \mathbb{E}[\log(1-v_j)]. \tag{25}$$

The expectations in Eq. 25 evaluate to

$$\begin{aligned} \mathbb{E}_{q(v,A)}[\log|A_k|] &= \langle \log|A_k| \rangle = \sum_{j=1}^{p} \psi(\frac{\nu_k + 1 - j}{2}) + p\log 2 + \log|A_k| \\ \mathbb{E}_{q(v,A)}[\gamma_i^T A_k \gamma_i] &= \nu_k \mathrm{Tr}(\gamma_i \gamma_i^T L_k) \\ \mathbb{E}_{q(v,A)}[\log v_k] &= \langle \log v_k \rangle = \psi(a_k) - \psi(a_k + b_k) \\ \mathbb{E}_{q(v,A)}[\log(1-v_k)] &= \langle \log(1-v_k) \rangle = \psi(b_k) - \psi(a_k + b_k). \end{aligned} \tag{26}$$

where $\psi(x) = \frac{d}{dx}\log(\mathit{\Gamma}(x)) = \frac{\mathit{\Gamma}'(x)}{\mathit{\Gamma}(x)}$ is the digamma function.

The updated $q(z)$ follows a Categorical distribution

$$q(z) = \prod_{i=1}^{J}\prod_{k=1}^{K} r_{ik}^{z_{ik}}, \quad \text{with } r_{ik} = \frac{\rho_{ik}}{\sum_j \rho_{ij}}. \tag{27}$$

*Variational M-step.* Next, we need to update $q(v)$ and $q(A)$ in each iteration.

$$
\begin{aligned}
\log q(v, A) &= \mathbb{E}_{q(z)}[\log p(\gamma, z, v, A)] + C \\
&= \mathbb{E}\left[\log \prod_{i=1}^{J}\prod_{k=1}^{\infty}\left[\mathcal{N}(\gamma_i|0, A_k^{-1})\right]^{z_{ik}}\right] + \mathbb{E}\left[\log \prod_{i=1}^{J}\mathrm{Cat}(z_i|\pi)\right] \\
&\quad + \mathbb{E}\left[\log \prod_{k=1}^{\infty}\mathrm{Beta}(v_k|a_0, b_0)\right] + \mathbb{E}\left[\log \prod_{k=1}^{\infty}\mathcal{W}(A_k|L_0, \nu_0)\right] + C \\
&= \sum_{i=1}^{J}\sum_{k=1}^{K} r_{ik}\left(\frac{1}{2}\log|A_k| - \frac{1}{2}\gamma_i^T A_k \gamma_i\right) + \sum_{i=1}^{J}\sum_{k=1}^{K}\mathbb{E}\left[z_{ik}\log \pi_k\right] \\
&\quad + \sum_{k=1}^{K}\left[(a_0 - 1)\log v_k + (b_0 - 1)\log(1 - v_k)\right] \\
&\quad + \sum_{k=1}^{K}\left[\frac{\nu_0 - p - 1}{2}\log|A_k| - \frac{1}{2}\mathrm{Tr}(L_0^{-1}A_k)\right] + C,
\end{aligned}
\tag{28}
$$

where $\sum_{k=1}^{K}\mathbb{E}\left[z_{ik}\log \pi_k\right] = \sum_{k=1}^{K}\left[\sum_{j=k+1}^{K} r_{ij}\log(1 - v_k)\right] + r_{ik}\log v_k$.

Separating the expression into terms that depend only on $v$ and those that only depend on $A$, the approximate posterior becomes

$$
\log q(v, A) = \log q(v) + \log q(A). \tag{29}
$$

For $v$ terms:

$$
\begin{aligned}
\log q(v) &= \sum_{i=1}^{J}\sum_{k=1}^{K}\left[\left(\sum_{j=k+1}^{K} r_{ij}\log(1 - v_k)\right) + r_{ik}\log v_k\right] x \\
&\quad + \sum_{k=1}^{K}\left[(a_0 - 1)\log v_k + (b_0 - 1)\log(1 - v_k)\right] + C \\
&= \sum_{k=1}^{K}\left(a_0 - 1 + \sum_{i=1}^{J} r_{ik}\right)\log v_k + \left(b_0 - 1 + \sum_{i=1}^{J}\sum_{j=k+1}^{K} r_{ij}\right)\log(1 - v_k) + C.
\end{aligned}
\tag{30}
$$

Therefore, $q(v)$ is a product of Beta distributions $q(v) \propto \prod_{k=1}^{K-1}\mathrm{Beta}(v_k|a_k, b_k)$, where the product stops at $K - 1$ as $q(v_k = 1) = 1$ by the truncated stick-breaking construction, and

$$
a_k = a_0 + \sum_{i=1}^{J} r_{ik},\ b_k = b_0 + \sum_{i=1}^{J}\sum_{j=k+1}^{K} r_{ij}. \tag{31}
$$

For $A$ terms:

$$
\begin{aligned}
\log q(A) &= \sum_{i=1}^{J}\sum_{k=1}^{K} r_{ik}\left(\frac{1}{2}\log|A_k| - \frac{1}{2}\gamma_i^T A_k \gamma_i\right) \\
&\quad + \sum_{k=1}^{K}\left[\frac{\nu_0 - p - 1}{2}\log|A_k| - \frac{1}{2}\mathrm{Tr}(L_0^{-1}A_k)\right] + C.
\end{aligned}
\tag{32}
$$

Match the terms with the product of Wishart distributions

$$
\begin{aligned}
\log q(A) &= \sum_{k=1}^{K}\log \mathcal{W}(A_k|L_k, \nu_k) \\
&= \sum_{k=1}^{K}\log W(L_v, \nu_k) + \frac{\nu_k - p - 1}{2}\log|A_k| - \frac{1}{2}\mathrm{Tr}(L_k^{-1}A_k),
\end{aligned}
\tag{33}
$$

and use the following property

$$x^T A x = \mathrm{Tr}(x x^T A),$$

we can write $q(A)$ as a product of Wishart distributions $q(A) \propto \prod_{k=1}^{K} \mathcal{W}(A_k|L_k, \nu_k)$ with

$$L_k = \left( L_0^{-1} + \sum_{i=1}^{J} r_{ik} \gamma_i \gamma_i^T \right)^{-1}, \; \nu_k = \nu_0 + \sum_{i=1}^{J} r_{ik}. \tag{34}$$

*Convergence check.* The convergence of the optimization can be checked by evaluating the ELBO after each iteration of the E and M-steps:

$$\begin{aligned} \mathrm{ELBO(q)} &= \sum_z \int_v \int_A q(z, v, A) \log \frac{p(\gamma, z, v, A)}{q(z, v, A)} \, dv \, dA \\ &= \mathbb{E}[\log p(\gamma|z, A)] + \mathbb{E}[\log p(z|\pi)] + \mathbb{E}[\log p(v)] + \mathbb{E}[\log p(A)] \\ &\quad - \mathbb{E}[\log q(z)] - \mathbb{E}[\log q(v)] - \mathbb{E}[\log q(A)], \end{aligned} \tag{35}$$

where $\mathbb{E}[\cdot]$ is with respect to $q(z, v, A)$. Each term in Eq. 35 can be evaluated separately; here we do not exhaustively enumerate the evaluations.

*Handling of local optimum.* VI is known to converge to a local maximum. We therefore choose to run the inference procedure multiple times, each starting from a different initialization, and aggregate the multiple runs to reduce the chance of being trapped in sub-optimal clustering. This is done by using the majority $K^*$ value from the runs and choosing the run that yields a model with lowest Bayesian information criterion (BIC) across all runs with the $K^*$ value.

##### A.9 Univariate gene enrichment test

The (univariate) gene-level test statistic is calculated as

$$\tilde{Q}_g = \tilde{\beta}_g^T \tilde{\beta}_g, \tag{36}$$

where the subscript $\cdot_g$ indicates all variants in the gene $g$. According to [14], such a quadratic form of SNP effects is commonly used in gene enrichment methods. Based on the SNP-level null hypothesis $H_0 : \mathbb{E}[\beta_i^2] \leq \sigma_\epsilon^2$, $\tilde{Q}_g$ is tested against the gene-level enrichment null hypothesis $H_0 : Q_g = 0$ that is dependent on $\sigma_\epsilon^2$. The normality assumption for true effects allow a linear combination of chi-squared test statistics to be used to test the significance of the gene. *gene-$\epsilon$* uses Imhof's method [24], which is implemented in `R`. Alternatively, tests like [19] and [28] can also be used. In this study, to compare the univariate gene-level results using different shrinkage methods, we implement the gene enrichment test in `Python` using [28] available through the package `chiscore`. We correct the output p-values for multiple testing by controlling the controls the false discovery rates (FDR) at level $\alpha = 0.05$ [4].

*Verification with biological processes of genes.* We use the gene annotation of SNPs from NCBI's Reference Sequence (RefSeq) database [36] in the UCSC Genome Browser, same as in [14]. The Gene Ontology (GO) knowledge base [3,2] provides a good resource to assess the gene enrichment test results. It contains knowledge about the biological processes that gene products may carry out, with which we can investigate the functions of genes that are identified as associated, possibly to varying degrees, to the traits of interest. We use the gene set enrichment analysis tool Enrichr [13] to verify identified genes against their known GO biological processes (Fig. 3C and D).

##### A.10 Categorizations of the associations types of clusters

Variants in each clusters can roughly be grouped into three types: trait-specific, shared-association, and spuriously associated. In the bivariate Gaussian case, each cluster has a variance-covariance matrix $\Sigma_k$. We compute the ratio between the largest and smallest eigenvalues of $\Sigma_k$. We also look the angle between x- or y-axis and the vector represented by the eigenvector corresponding to the largest eigenvalue, which represents the direction of the major axis of the confidence ellipse of the Gaussian. If the ratio is large enough (e.g., $> 20$) and the angle is small (e.g., $< 15°$ to either axis), then we designate the cluster to be trait-specific.

If the cluster is not trait-specific, then we categorize it to have shared association. However, this type of clusters may sometimes contain variants that have large trait-specific effects, but get mixed together with those having large effects in both traits. We are grouping all these as "shared association", although finer-scaled categorization is possible. For clusters that have $\Sigma_k$ with both eigenvalues small, i.e., having a small $\mathrm{Tr}(\Sigma_k)$, we treat them as spuriously associated. The threshold for $\mathrm{Tr}(\Sigma_k)$ can be adapted based on observed patterns in the output.

##### A.11 Comparison to a model capturing only linear relationships

One advantage of using neural-network-based effect size shrinkage is that, unlike elastic net which assumes linear relationships between true and inflated effects, network architecture design like the *ML-MAGES 2L and 3L* can easily capture non-linearities among correlated effects of variants.

To assess the necessity of capturing non-linear relationships, we further evaluate the performance of ML-MAGES framework using a single-layer neural network that directly connects all the input features to the output. The size of this single layer is determined by the number of features. This special network is only capable of capturing linear relationships, thus serves as a baseline comparison for showing how important non-linearity is in addressing effect size inflation. The performance of this special network is labeled as Linear in Fig. 7 and Fig. 8 in the upcoming sections. It is not surprising that *ML-MAGES (2L) and ML-MAGES (3L)* consistently outperform *Linear*, which demonstrates the importance of capturing non-linear relationships among correlated effects in effect size shrinkage. The precision-recall performance of *Linear* is worse than that of elastic net shrinkage, which is expected given that elastic net shrinkage incorporates effects of all variants in the segment as inputs. In contrast, *Linear* adheres to the *ML-MAGES* framework by focusing solely on a selected set of variants with the highest correlation to the focal variant.

##### A.12 Additional simulation comparisons in a multi-trait scenario

To show that our *ML-MAGES* framework is generally applicable to multi-trait association patterns, in addition to the performance comparison we conducted in Section 3.1 with two simulated traits, we extended our simulation to a three-trait scenario. The generation of simulation data closely follows that in Section 3.1 and Appendix A.4, and is detailed below.

A total of 100 simulations are generated. In each simulation, using a sub-sampled set of $N^{(s)} = 10000$ individuals from the UKB genotype data in chormosome 15, we randomly select 10% of genes to be truly associated to some traits, categorized into five association types: associated with only one trait (three types: trait 1-specific, trait 2-specific, and trait 3-specific), associated with two traits (traits 1&2-shared), or associated with all three (all traits-shared). To limit scenario complexity, we excluded certain pairwise combinations for shared associations between two traits. Since traits are synthetic, we generalized this scenario by focusing on traits 1 and 2 without loss of generality. Within each such gene, either 40% of variants or 2 variants—whichever is larger—are simulated to be associated, i.e., having non-zero true effects. True effects are initially sampled from zero-mean normal distributions with variance uniformly drawn from [0.1, 0.5]. For variants with shared associations across multiple traits, effects are generated from a zero-mean multivariate normal distribution. The covariance matrices for these distributions are selected from a pre-generated pool of 50 positive-definite symmetric matrices, each validated to ensure significant pairwise correlations ($p < 0.01$) among variables. The narrow-sense heritability of all traits was fixed at $h^2 = 0.8$. Because we simulated three traits, there are 300 single-trait simulations in total, and performance comparisons of shrinkage methods for all simulations are summarized in Fig. 7. Fig. 1 illustrates the performance of gene-level multivariate analysis for three of the five ground-truth association types: trait 1-specific, traits 1&2-shared, and all traits-shared. Trait 2- and trait 3-specific associations are omitted for brevity but yield results comparable to trait 1. In both SNP-level shrinkage and gene-level association pattern identification, our method *ML-MAGES* consistently shows good performance.

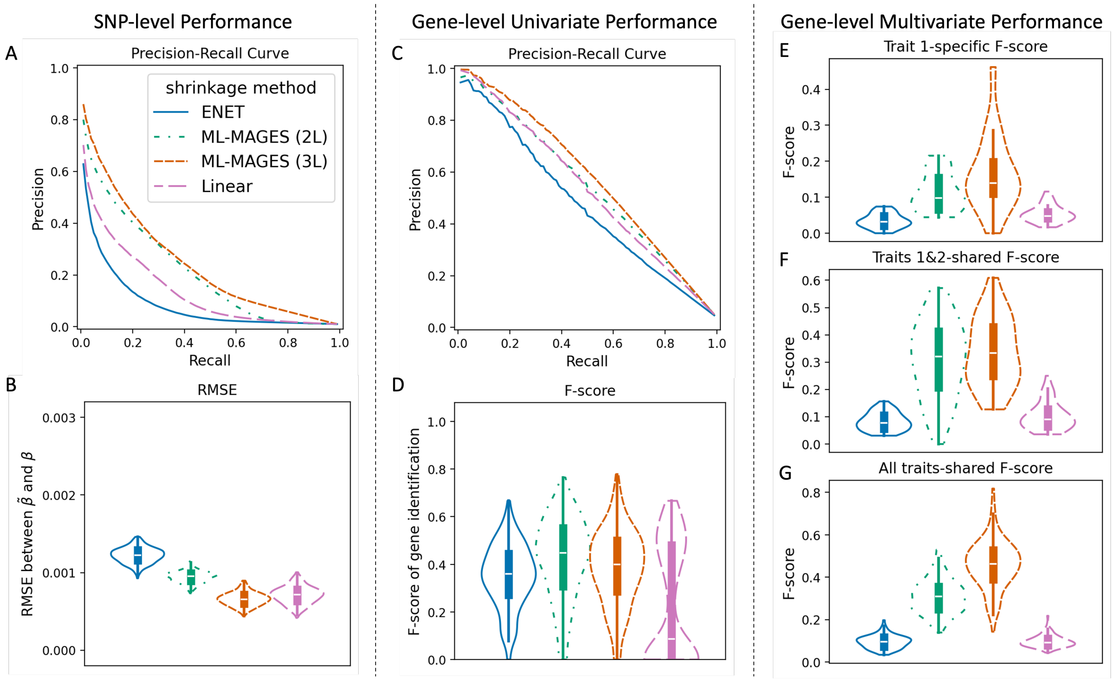


**Fig. 7. Our NN methods *ML-MAGES 2L* and *3L* outperform the others in shrinking inflated GWA effect sizes and subsequently identifying associated genes in simulations with three traits.** The figure design follows that of Fig. 2 in the main text. Legends shown in panel A apply to all panels; each violin plot ordered from left to right as *enet*, *ML-MAGES 2L* and *3L*, and *Linear*. **Left column**: SNP-level performance, evaluated by comparing the regularized effects and the true effects of each simulation. **Middle column**: gene-level single-trait association performance analysis, evaluated by comparing univariate enrichment test with the simulated ground truth. **Right column**: gene-level multi-trait association performance analysis, evaluated by comparing multivariate clustering output with the simulated ground truth of association types (i.e., trait-specific vs. shared). **A**: Precision-recall curve (PRC) averaged across all 100 simulations (by interpolation), where the positives are the true non-zero effects and the precision-recall pairs are obtained by thresholding $|\tilde{\beta}|$. **B**: RMSE between $\beta$ and $\tilde{\beta}$. **C**: PRC of each method averaged across all simulations (by interpolation), where the true positives are the true associated genes and the precision-recall pairs are obtained for different p-values of gene enrichment tests. **D**: F-score of identifying true associated genes, where genes with a FDR-adjusted $p < 0.05$ from the enrichment analysis is identified as associated. **E**: Trait-specific F-scores for identifying genes only associated to the synthetic trait 1, when ranking the genes by their fraction of variants in trait-specific clusters and compared against the ground-truth. **F**: F-score for identifying genes associated to both traits 1 and 2, when ranking the genes by their fraction of variants in clusters of shared association between these two traits and compared against the ground-truth. **G**: F-score for identifying genes associated to all three traits 1, 2, and 3, when ranking the genes by their fraction of variants in clusters of shared association among all traits and compared against the ground-truth.

#### A.13 Additional simulation comparisons when training uses imputation data

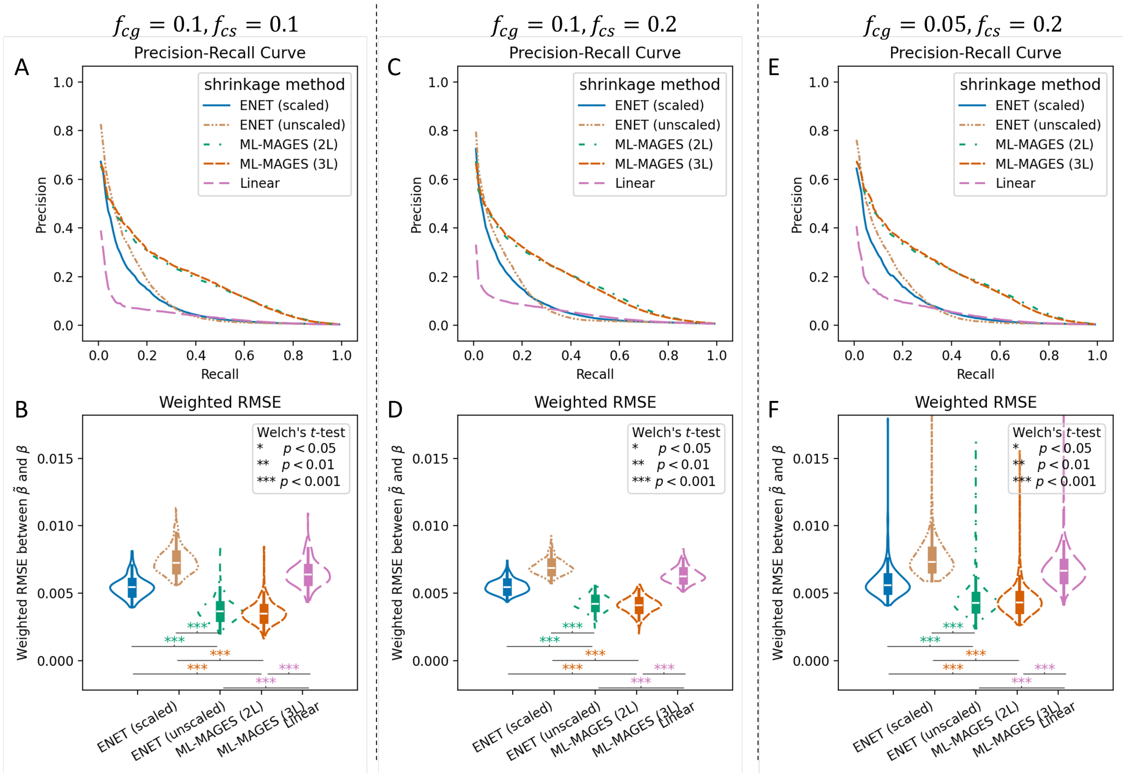


**Fig. 8.** ***ML-MAGES 2L and 3L* outperform elastic net and linear neural network model on shrinking inflated GWA effect sizes when models are trained on synthetic data simulated from imputed UKB genotypes.** Outputs of 10 separately trained models for each NN architecture are ensembled to provide the final shrinkage results. The three simulation settings, **A-B**, **C-D**, and **E-F**, differ in the fraction of causal genes ($f_{cg}$) and the fraction of causal variants in each causal gene ($f_{cs}$), labeled on top of each column). The fraction of causal variants no in any causal genes is fixed to be 0.1. The two NN architectures, *ML-MAGES* (2L) and *ML-MAGES* (3L), and the single-layer neural network for comparison, labeled as *Linear*, are each averaged across 10 models trained independently. The performances of *ML-MAGES* (2L) and *ML-MAGES* (3L) are compared to that of *Linear*, as well as that of elastic net using both untransformed and transformed synthetic effects, labeled as "scaled" and "unscaled". The transformed synthetic effects are scaled to match the distribution of the summary statistics of mean corpuscular volume (MCV) from UKB and are used to construct NN inputs. The unscaled effects closely reflect the simulation that [14] used for evaluating gene-e. The figure style follows that of Fig. 2 panel A and B, where on top shows the precision-recall curves and on bottom shows the weighted RMSE, with the weights applied inversely proportional to the fraction of variants with true non-zero and zero effects. The significance of the comparisons using Welch's t-test are indicated on the bottom of the violin plots.

To further demonstrate the reliability of the method, we train the models using simulation based on imputed genotype data of UKB, which contains regions with much denser variants subjecting to much higher correlations than the genotyped data, and use ensemble learning results for effect size shrinkage. We then apply them to simulated evaluation data based on genotyped data of chromosome 15 (15,250 SNPs), same as before. The results are shown in Fig. 8.

The training simulation is generated based on imputation data from chromosomes 7 to 22, which contain a total of 2,283,694 variants (the other chromosomes are not included for computational burden caused by processing large number of variants in imputation data). Simulated samples based on chromosomes 10, 13, 17, and 19 are used in the testing set and the rest in the training set. The difference in data density leads to significant variation in the correlations between variants. Highly-correlated markers are particularly subject to inflation, making it difficult to shrink spurious effect sizes, as these spurious effects can easily surpass true effects due to high LD. As a result, models trained on imputed data cannot fully reduce spurious associations to close-to-zero values while retaining the effects of putative true associations (results not shown). To improve the robustness of the shrinkage method, we incorporate ensemble learning, where we aggregate the results from ten independently trained models for each architecture by averaging the regularized effects from all ten models. To account for non-sparsity caused by marker density, we also measure the performance via both PRC and weighted RMSE, where the weights are applied inversely proportional to the fraction of variants with true non-zero and zero effects.

In simulation, we vary the fraction of causal genes ($f_{cg}$) and the fraction of causal variants in each causal gene ($f_{cs}$), while fixing the fraction of causal variants not in any causal genes to be 0.1. The simulated effects are then transformed, as described in Appendix A.4, to match the distribution of estimated GWA effects of the trait mean corpuscular volume (MCV) in the imputation data. We choose this trait that has not been used in our previous analyses to demonstrate the generalizability of performance evaluations. Each simulated data sample is based on a chromosomal segment of 1,000 variants from the imputation data. We generate 150 such data samples for each of the chromosome, resulting in a total of 600,000 data samples from training chromosomes and 200,000 from validation chromosomes. From them, a total of 15,000 are subset and used for training, and 5,000 are used for validation. When effect values are very small, training can suffer from numerical instability of some weights or gradients in the neural networks; therefore, we scaled the simulated effects by 250, which equals one over the absolute value of the original transformed effects rounded to the nearest 10. Note that when applying the trained models, inputs are scaled up by 250 as well, and the corresponding outputs are then divided by 250 afterwards.

We evaluated the model's performance on simulation data generated from genotyped data. The difference between the simulation of training and evaluation data, as well as the difference in simulation settings, allow us to demonstrate robustness of *ML-MAGES* when the training simulations do not perfectly reflect the true underlying effect size distribution, which is often the case in practice. We also include the performance of elastic net shrinkage using untransformed synthetic effects, as the original elastic net method does not depend on the generation or transformation of training data. Our NN-based shrinkage consistently outperforms elastic net shrinkage, both with and without effect transformation, as shown in Fig. 8.

##### A.14 Implementation details

We implement the method and perform our analyses in `Python 3.9`. Neural networks are implemented using the package `PyTorch`. Elastic net shrinkage is implemented using the package `scikit-learn`. *SuSiE-RSS* (*susie*) is performed in `R` using the library `susieR` [56,64]. We conducted our non-training tasks on a machine with Linux 5.14 OS, x86-64 architecture, Intel Xeon Processor E5-2675, using 1 CPU node. Model training was carried out on 1 GPU and 1 CPU node featuring NVIDIA GeForce RTX 3090, taking less than an hour for each model (Table 6). Experiments shown in our study require memory of no more than 128G per which can be split and parallelized, e.g., on a single chromosome.

The implementation is available at `https://github.com/ramachandran-lab/ML-MAGES`. We also provide example data to run our method in the repository.

#### B Supplementary Tables

We include here Tables 1-6 that complement the performance comparison results of 8 shrinkage methods—*enet*, *susie*, and 6 NN-based methods—shown in Figs. 2 and 3. Measures are shown with their 95%-CI over all simulations. Labeling of the methods is different from Figs. 2 and 3: *fc2top5* denotes the NN model using two fully-connected layers and top $T = 5$ variants for the features and so forth. So *ML-MAGES (2L)* in the main text is now labeled as *fc2top15*, and *ML-MAGES (3L)* becomes *fc3top15*.

**Table 1. SNP-level performance of the 8 shrinkage methods (*enet*, *susie*, and 6 neural-network models) on SNP-only simulations.** The average measures over 200 simulations are reported, followed by their 95%-CI in brackets. Average precision is the area under the precision-recall curve (PCR). Best value for each measure is bolded.

| Method | RMSE ($\times 10^2$) | Pearson Correlation | Average Precision |
|---|---|---|---|
| enet | 0.135 [0.129, 0.140] | 0.349 [0.338, 0.360] | 0.194 [0.176, 0.213] |
| susie | 0.147 [0.120, 0.174] | 0.561 [0.534, 0.589] | 0.282 [0.253, 0.310] |
| fc2top5 | 0.098 [0.093, 0.104] | 0.549 [0.527, 0.570] | 0.282 [0.257, 0.308] |
| fc2top10 | 0.098 [0.092, 0.103] | 0.538 [0.512, 0.564] | 0.272 [0.245, 0.299] |
| fc2top15 | **0.096** [0.090, 0.101] | 0.572 [0.550, 0.595] | 0.286 [0.261, 0.310] |
| fc3top5 | 0.103 [0.097, 0.108] | 0.564 [0.535, 0.592] | 0.254 [0.228, 0.280] |
| fc3top10 | 0.098 [0.093, 0.103] | 0.573 [0.544, 0.603] | 0.286 [0.260, 0.312] |
| fc3top15 | 0.098 [0.093, 0.103] | **0.623** [0.601, 0.645] | **0.287** [0.260, 0.314] |

**Table 2. SNP-level performances of the 8 shrinkage methods (same as in Table 1) on gene-level simulations.** The average measures over 200 simulations are reported, followed by their 95%-CI in brackets. Average precision is the area under the PRC shown in Fig. 2A. RMSE are shown in Fig. 2B. Best value for each measure is bolded.

| Method | RMSE ($\times 10^2$) | Pearson Correlation | Average Precision |
|---|---|---|---|
| enet | 0.118 [0.117, 0.120] | 0.142 [0.135, 0.148] | 0.086 [0.079, 0.093] |
| susie | 0.219 [0.215, 0.223] | **0.457** [0.440, 0.475] | **0.257** [0.245, 0.270] |
| fc2top5 | 0.057 [0.056, 0.059] | 0.397 [0.387, 0.407] | 0.232 [0.222, 0.242] |
| fc2top10 | 0.065 [0.063, 0.066] | 0.405 [0.395, 0.415] | 0.222 [0.212, 0.231] |
| fc2top15 | 0.071 [0.070, 0.073] | 0.398 [0.388, 0.409] | 0.219 [0.210, 0.229] |
| fc3top5 | **0.047** [0.046, 0.048] | 0.443 [0.430, 0.456] | 0.225 [0.215, 0.235] |
| fc3top10 | 0.051 [0.049, 0.052] | 0.440 [0.427, 0.452] | 0.230 [0.220, 0.240] |
| fc3top15 | 0.049 [0.048, 0.050] | 0.455 [0.443, 0.467] | 0.228 [0.219, 0.238] |

**Table 3. Performance of gene enrichment analysis of the 8 shrinkage methods (same as in Table 1) on gene-level simulations.** The average measures over 200 simulations are reported, followed by their 95%-CI in brackets. F-scores are shown in Fig. 2D. Best value for each measure is bolded.

| | Precision | Recall | F-score |
|---|---|---|---|
| enet | 0.443 [0.402, 0.483] | 0.456 [0.425, 0.488] | 0.355 [0.334, 0.375] |
| susie | 0.309 [0.289, 0.329] | **0.832** [0.813, 0.851] | 0.428 [0.409, 0.447] |
| fc2top5 | 0.701 [0.670, 0.733] | 0.428 [0.404, 0.453] | 0.494 [0.473, 0.514] |
| fc2top10 | 0.649 [0.615, 0.683] | 0.479 [0.450, 0.508] | **0.498** [0.478, 0.519] |
| fc2top15 | 0.583 [0.550, 0.616] | 0.497 [0.469, 0.526] | 0.483 [0.464, 0.502] |
| fc3top5 | 0.662 [0.623, 0.700] | 0.473 [0.442, 0.504] | 0.475 [0.456, 0.494] |
| fc3top10 | 0.731 [0.697, 0.765] | 0.413 [0.388, 0.438] | 0.490 [0.468, 0.513] |
| fc3top15 | **0.822** [0.792, 0.853] | 0.351 [0.328, 0.374] | 0.464 [0.442, 0.486] |

**Table 4. Performances of gene-level bivariate analysis—trait-specific versus shared-association categorization—of the 8 shrinkage methods (same as in Table 1) on gene-level simulations.** The average measures over 200 simulations are reported, followed by their 95%CI in brackets. Measures are those shown in Fig. 2E-G. Best value for each measure is bolded.

| | Trait-1-specific F-score | Trait-2-specific F-score | Shared-association F-score |
|---|---|---|---|
| enet | 0.095 [0.095, 0.095] | 0.101 [0.093, 0.108] | 0.095 [0.089, 0.101] |
| susie | 0.093 [0.082, 0.103] | 0.087 [0.078, 0.096] | 0.085 [0.077, 0.092] |
| fc2top5 | 0.312 [0.284, 0.341] | **0.378** [0.342, 0.413] | 0.404 [0.387, 0.420] |
| fc2top10 | 0.126 [0.109, 0.143] | 0.166 [0.144, 0.189] | 0.398 [0.381, 0.416] |
| fc2top15 | **0.325** [0.312, 0.339] | 0.322 [0.307, 0.337] | 0.408 [0.392, 0.425] |
| fc3top5 | 0.236 [0.214, 0.259] | 0.261 [0.237, 0.285] | 0.267 [0.244, 0.291] |
| fc3top10 | 0.270 [0.252, 0.289] | 0.229 [0.209, 0.249] | **0.458** [0.438, 0.477] |
| fc3top15 | 0.273 [0.257, 0.289] | 0.246 [0.222, 0.269] | 0.407 [0.389, 0.425] |

**Table 5. Shrinkage time (in seconds) of the 8 shrinkage methods (same as in Table 1) for each simulation in the gene-level simulation data.** Results are calculated across 200 simulations generated based on chromosome 15 (15,250 SNPs) splitted into 17 LD blocks. *SuSiE-RSS* is run in `R`, while the rest are run in `Python`. Implementation details can be found in Appendix A.14.

| | enet | susie | fc2top5 | fc2top10 | fc2top15 | fc3top5 | fc3top10 | fc3top15 |
|---|---|---|---|---|---|---|---|---|
| Median | 15.29 | 31.83 | 0.84 | 0.84 | 0.84 | 0.84 | 0.84 | 0.84 |
| Mean | 15.88 | 33.62 | 0.86 | 0.86 | 0.86 | 0.86 | 0.86 | 0.86 |
| Std.Dev. | 2.99 | 8.72 | 0.06 | 0.07 | 0.08 | 0.09 | 0.08 | 0.08 |

**Table 6. Training time (in seconds) per epoch for each of the 6 NN models.** Median, mean, and standard deviation of the time spent per epoch during the training of each model are shown. The total number of epochs each model takes until training finishes is also included.

| | fc2top5 | fc2top10 | fc2top15 | fc3top5 | fc3top10 | fc3top15 |
|---|---|---|---|---|---|---|
| Median | 15.36 | 15.48 | 20.18 | 19.91 | 25.11 | 18.98 |
| Mean | 15.38 | 15.49 | 19.80 | 19.92 | 24.87 | 18.98 |
| Std.Dev. | 0.08 | 0.05 | 0.70 | 0.05 | 0.83 | 0.86 |
| Number of epochs | 28 | 47 | 47 | 60 | 81 | 68 |
